# An adaptive multi-strategy metaheuristic for robust model calibration in large-scale systems biology

**DOI:** 10.64898/2026.01.27.701977

**Authors:** Andrea Polo-Rodríguez, David R. Penas, Julio R. Banga

**Affiliations:** Computational Biology Lab, MBG-CSIC (Spanish National Research Council), Pontevedra, Galicia, Spain; Applied Mathematics Dept., Universidade de Santiago de Compostela, Santiago de Compostela, Galicia, Spain

**Keywords:** Hybrid metaheuristic, global optimization, model calibration, dynamic modelling, systems biology, computational biology

## Abstract

Parameter estimation is a central challenge in systems biology, particularly for large dynamic models described by nonlinear ordinary differential equations (ODEs). These global optimization problems exhibit landscapes which are topologically heterogeneous, often exhibiting a pathological mixture of stiff, smooth valleys and rugged, noisy plateaus, making single-strategy hybrids ineffective. While methods like enhanced Scatter Search (`eSS`) represent the current state-of-the-art, their rigid intensification strategies can limit performance in large-scale or ill-conditioned scenarios. In this work, we introduce `eLSHADE+`, a novel metaheuristic architecture designed to adapt to these topological challenges. The proposed algorithm augments a recent Differential Evolution variant (`LSHADE`) with a probabilistic multistrategy hybridization. Unlike standard memetic algorithms, `eLSHADE+` uses three distinct operational modes: (i) gradient-based intensification for precision in differentiable regions, (ii) derivative-free search for robustness against numerical noise, and (iii) pure global exploration to conserve computational resources in complicated basins. Additionally, a logarithmic parameter space transformation is incorporated to facilitate search across multi-scale biological constants. We rigorously evaluated `eLSHADE+` using the BioPreDyn benchmark suite, comprising challenging real-world problems. Comparative analysis demonstrates that this adaptive multi-strategy approach yields statistically superior convergence speed and solution accuracy compared to `eSS` and other competitive metaheuristics, establishing a new baseline for robust model calibration in computational biology. Code and data are available at https://doi.org/10.5281/zenodo.18379327.

## 1. Introduction

Systems biology is an interdisciplinary field that integrates biology, mathematical modeling, and advanced computational techniques [1, 2]. Biological processes in nature are often characterized by high complexity, making them challenging to describe with qualitative models alone. Such models frequently fall short in capturing the intricate interconnections among system components. In contrast, mathematical models provide a framework for developing quantitative representations that can overcome these limitations. The incorporation of mathematical and computational approaches into the study of biological systems has significantly transformed the way these systems are understood.

Over the past few decades, systems biology has become a cornerstone of biological research, offering powerful tools for investigating complex biological processes. Systems biology has critical applications across major scientific fields. In agriculture, it is used to model gene networks to improve crop resilience and yield [3, 4]. In medicine, the approach clarifies complex diseases to identify novel drug targets and enable personalized therapies [5, 6]. Furthermore, in industrial biotechnology, it drives the rational engineering of microbial systems to optimize the production of biofuels, pharmaceuticals, and other chemicals [7].

Among the various modeling approaches available [8], models based on ordinary differential equations (ODEs) are particularly well-suited for describing and predicting the temporal dynamics of biological systems [1, 9]. These models incorporate parameters, which are quantities that may vary under different conditions but remain constant once specified. In other words, these parameters are the specific numerical values that tailor the model to a particular biological system. They represent real, physical quantities such as biochemical reaction rates and protein binding affinities.

In general, the values of most parameters are not directly observable from experimental data. Estimating these parameters by fitting the model to experimental data (e.g., time-course measurements of protein concentrations) is the crucial step that connects our abstract mathematical equations to real-world measurements [10– 12]. Without this, the model has no descriptive power. Once a model is calibrated with estimated parameters that allow it to accurately reproduce existing data, it becomes a predictive tool. We can then use it to perform *in silico* (computer-based) experiments that would be difficult, expensive, or impossible to do in a lab.

Parameter estimation, also known as model calibration, involves determining the values of unknown parameters that minimize the discrepancies between model predictions and observed data. This process can be formally framed as an optimization problem, where the objective function to be minimized quantifies the deviation between predictions and measurements. Due to the inherent complexity of biological systems, parameter estimation poses several significant challenges [11, 13, 14]. First, models often contain a large number of parameters, leading to high-dimensional problems. Second, the non-linear nature of most biological models typically results in non-convex objective functions, making the optimization landscape multimodal and prone to numerous local minima. Third, evaluating the objective function often requires computationally intensive simulations, further complicating the estimation process. Additional difficulties arise from limitations in the available data, which may be sparse or noisy. These challenges can lead to identifiability issues (either structural or practical) that compromise the reliability of parameter estimates and, consequently, the predictive power of the model [9, 11].

Here we consider parameter estimation in dynamic models of biological systems described by large sets of nonlinear ordinary differential equations (ODEs). Given time-series data, the task is to infer kinetic and regulatory parameters by minimizing a simulation-based objective function subject to ODE constraints. Due to the characteristics mentioned above, these problems are high-dimensional, nonlinear, often stiff, and computationally expensive, yielding rugged, multimodal landscapes. This motivates the use of global optimization methods that can robustly escape local minima while handling large scale, strong nonlinearity, and costly function evaluations. Effective solvers must balance exploration with judicious use of expensive simulations and exploit problem structure (constraints, bounds, sensitivities).

Global optimization methods can be classified, roughly, into deterministic and stochastic (probabilistic) approaches [15–18]. Deterministic methods systematically partition the search space and propagate bounds/relaxations to provide certificates or guarantees of global optimality, but tend to scale poorly with dimension and nonlinearity. Stochastic methods employ randomized exploration (e.g., evolutionary algorithms, simulated annealing, particle swarms, differential evolution, and other metaheuristics), trading formal guarantees for robustness and scalability on rugged objective landscapes.

Early studies established that stochastic global optimization could solve challenging ODE parameter estimation problems in systems biology [19], outperforming deterministic methods. However, their computational cost spurred the development of hybrid (memetic) metaheuristics, which combine a global search with local refinement and are now state-of-the-art (a more detailed review is provided in the next section). Among these, enhanced Scatter Search (`eSS`) [20] has consistently proven a top performer in benchmarks [21, 22]. Nevertheless, its effectiveness is limited by the high computational cost of repeated simulations and local searches, particularly for large-scale or stiff models [23, 24].

Motivated by these observations, this contribution presents a novel hybrid metaheuristic designed to address the specific topological challenges of biological parameter estimation. While hybrid algorithms are well-established, standard approaches typically rely on a single local search strategy, either gradient-based or derivative-free. We argue that this single-mode intensification is insufficient for biological landscapes, which are characterized by a heterogeneous mixture of deep, smooth valleys (dictated by stiff dynamics) and rough, noisy plateaus (caused by numerical integration tolerances and lack of identifiability).

### 1.1. Main Contributions

This work makes three principal contributions to the field of parameter estimation in systems biology:

#### 1. Probabilistic multi-strategy hybridization framework

We introduce a novel adaptive hybridization scheme that probabilistically selects among three operational modes: gradient-based intensification, derivative-free robust search, and pure global exploration (no local search). Unlike conventional memetic algorithms that commit to a single local optimizer, this tri-modal approach dynamically adapts to the heterogeneous topology of biological objective landscapes, balancing computational efficiency with robustness against numerical noise and ill-conditioning.

#### 2. Characterization of when pure exploration outperforms intensification

Through systematic ablation studies and analysis of convergence dynamics across diverse problem scales, we demonstrate that deliberately skipping local search in certain iterations, contrary to conventional hybrid design, can preserve computational budget for global exploration in rugged, deceptive regions of the search space. This finding challenges the assumption that more frequent local refinement always improves performance, particularly for stiff ODE systems and high-dimensional problems with expensive function evaluations.

#### 3. New state-of-the-art performance on biological benchmarks

We provide the first rigorous demonstration that LSHADE-based hybrid metaheuristics can systematically outperform the previous state-of-the-art (eSS) on realistic systems biology parameter estimation problems. Our comprehensive evaluation on a challenging benchmark suite establishes `eLSHADE+` as a new baseline method, achieving statistically significant improvements in solution quality (up to 2 orders of magnitude error reduction on some problems) and robustness (lower variance across independent runs) while maintaining comparable or superior computational efficiency.

Beyond these algorithmic contributions, we release MATLAB code providing a unified interface for comparing state-of-the-art global optimization methods on ODE-constrained problems, facilitating future research in this domain [25].

The remainder of this paper is organized as follows: Section 2 provides a detailed literature review with an assessment of the state of the art. Section 3 defines mathematically the parameter estimation problem, presents the proposed `eLSHADE+` algorithm (addressing contribution 1), discusses other recent competitive methods for comparison, and outlines the experimental methodology and benchmark suite. Section 4 presents our empirical findings (contributions 2–3), which are subsequently discussed in depth in Section 5. Finally, Section 6 summarizes the key contributions and considers their broader implications.

## 2. Literature survey

Since parameter estimation is inherently a minimization problem, its solution requires the use of an optimization algorithm. Optimization methods can be broadly categorized as either local or global, depending on their ability to find global or only local optima [26]. Local optimization algorithms can further be distinguished by the type of information they use about the objective function. Specifically, derivative-free methods rely solely on the function’s value, whereas gradient-based methods also incorporate information about its first derivatives or second-order derivatives, such as the Hessian matrix [26, 27]. Although parameter estimation in dynamic models is often tackled with local optimization algorithms [14], because the associated objective landscapes are typically nonconvex, these methods are prone to converging to suboptimal local minima [13, 28–30].

A common practical remedy is multi-start local search [31–33], in which a local optimizer is repeatedly initialized at diverse random points within the admissible parameter bounds. This multi-start approach is most successful for problems with favorable optimization landscapes characterized by a limited number of local minima and a large basin of attraction for the global optimum. It is particularly effective in low-to-moderate dimensional parameter spaces, especially when tight prior bounds reduce the search volume, thereby increasing the probability that a random start will lead to the global solution. Conversely, the strategy’s reliability diminishes significantly for more challenging problems. Its primary limitation is the curse of dimensionality, where the number of starts required to adequately cover a highdimensional space becomes computationally infeasible. Furthermore, the presence of rugged or deceptive landscapes with numerous local minima, strong parameter correlations creating narrow and elongated valleys, or numerically stiff ODE systems can severely impede the performance of the underlying local optimizer, often leading to premature convergence or a false sense of security in a suboptimal solution.

These limitations motivate the use of global optimization. When the objective landscape is multi-modal or contains extended flat valleys, the basin of attraction of the global solution can occupy only a tiny fraction of the feasible set, so that multi-start local search becomes impractical. Global optimization algorithms can be classified into two main categories: deterministic methods, which guarantee convergence to a global solution within finite time, and stochastic methods (including metaheuristics), which do not offer such guarantees but are often more scalable and flexible [16, 34].

In parameter estimation, global optimization methods introduce a more principled exploration and bounding. Crucially, global optimization also serves model diagnostics by exposing competing optima due to weak identifiability, quantifying non-uniqueness (via level sets or profile likelihoods), and preventing conclusions driven by arbitrary local minima. Although more computationally demanding, many of these methods are highly parallelizable and, for challenging nonlinear ODE models, often the only reliable route to trustworthy parameter estimates and robust experimental design.

### 2.1. Deterministic global optimization

Deterministic global optimization methods are highly desirable because they are the only approaches that can provide a formal guarantee of finding the global optimum within a specified tolerance. Esposito and Floudas [28] pioneered a deterministic global optimization framework for ODE-constrained parameter estimation, using spatial branch-and-bound with convex relaxations to obtain certificates of global optimality and demonstrating its effectiveness on nonlinear kinetic models. Building on these early efforts, a substantial literature expanded and refined the use of deterministic global optimization in chemical kinetics, advancing spatial branch-and-bound, convex/interval relaxations, and validated integration for ODE-constrained estimation [35–41].

In parallel, analogous approaches were brought into systems biology, where several studies reported globally optimal (or certified near-global) parameter estimates for mechanistic dynamic models [42–45]. For recent comprehensive overviews of rigorous global optimization in this context, covering algorithms, verification, and open challenges, see [46, 47]. Notwithstanding these significant advancements, the practical scope of deterministic global optimization for dynamic systems remains limited, as highlighted in a recent review [47]. State-of-the-art methods struggle to scale beyond problems of a modest size, typically involving less than ten parameters and states. Consequently, while extending their reach is an active area of research [48–51], their utility for solving the high-dimensional models characteristic of many real-world biological systems is, at present, restricted.

### 2.2. Stochastic global optimization

Early studies in systems biology demonstrated that stochastic global optimization, including metaheuristics, could solve ODE parameter-estimation instances that deterministic global solvers and purely local methods could not handle within practical resources [19, 52, 53]. Already from these studies it was clear that pure stochastic methods were too computationally expensive for this class of problems.

Metaheuristics are a class of optimization methods that incorporate stochastic elements into the search process. Depending on their underlying strategies, metaheuristics can be broadly categorized into several families. These include evolutionary algorithms (such as genetic algorithms), simulated annealing, and other populationbased or trajectory-based approaches, such as ant colony optimization, tabu search, and particle swarm optimization. Each of these methods balances exploration and exploitation in different ways, making them particularly suitable for high-dimensional, multimodal, and computationally expensive problems [10, 34, 54].

Subsequent research has built upon these earlier studies by exploring a variety of stochastic global optimization methods. Sun and Garibaldi [55] provided a comprehensive review of metaheuristics applied to parameter estimation in systems biology. They also reviewed initial efforts making use of pure stochastic algorithms and simpler metaheuristics, such as Simulated Annealing, Genetic Algorithms, Evolution Strategies, and Differential Evolution, highlighting the versatility of these approaches in addressing the intricate challenges posed by biological modeling.

Notably, this review [55] further emphasized the significant advantages of hybrid metaheuristics over their pure counterparts and basic stochastic algorithms, particularly in enhancing effectiveness, efficiency, and robustness when dealing with complex problems. These hybrids excel in improving solution quality and reducing computational time, as demonstrated by other studies [56]. Overall, these results advocate for increased attention to hybridizing metaheuristics with classical algorithms to further boost their performance, asserting that such approaches are indispensable for navigating the noise, complexity, and computational demands of real-world biological systems.

Regarding these hybrid metaheuristics, Rodriguez-Fernandez et al. [57] introduced a hybrid method that merged `SRES`, an evolution strategy designed for global search, with the `DN2GB` local solver (a sophisticated algorithm combining Gauss-Newton and quasi-Newton techniques, enhanced by a trust region approach for stability). This approach markedly outperformed the earlier results of Moles et al. [19], which relied on various stochastic global optimization methods. Shortly thereafter, the same group developed `SSm`, a novel hybrid metaheuristic integrating scatter search with a suite of specialized local solvers [58, 59], demonstrating exceptional efficiency in solving complex problems and surpassing prior methods.

Building on `SSm`, Egea et al. [20, 60] introduced enhanced Scatter Search (`eSS`), an evolved version that simplifies yet strengthens the original design to address common hurdles in nonlinear dynamic system optimization, such as noise, flat regions, non-smoothness, and discontinuities. `eSS` strikes a robust balance between global exploration and efficiency, incorporating a local search procedure to expedite convergence to optimal solutions. Drawing on scatter search and path relinking, `eSS` employs a small, quality-selected population from diverse initial solutions and systematically combines them, while strategies based on path relinking generate new solutions biased toward the best population members. Parallel versions of enhanced Scatter Search (`eSS`) were developed [61] to address large-scale problems, enabling solutions within reasonable wall clock times.

A number of studies have shown the consistent success of `eSS` in benchmarks [21, 22, 62] and various challenging applications [23, 24, 45, 56, 63–77]. However, some of these more recent evaluations have also shown that the effectiveness of `eSS` for large-scale problems can be constrained by the high computational cost of repeated simulations and local searches. Although distributed computing can alleviate the computational burden associated with complex parameter estimation problems [24], even high-performance computing infrastructures often struggle to manage the most challenging cases within reasonable timeframes [23], highlighting areas for improvement by new methods.

These results are consistent with other areas which have also reported that the performance of hybrid metaheuristics often deteriorates in Large-Scale Global Optimization (LSGO) problems compared to their performance in lower-dimensional cases [78–80]. This limitation has driven extensive research over the past decades, as shown by numerous publications in leading journals focused on comparing the performance of various optimization metaheuristics and introducing increasingly challenging LSGO benchmark problems in the CEC competitions [81–85].

Here, we focus on the problem of parameter estimation in systems biology. To address these limitations, we aim to develop a novel hybrid metaheuristic that is able to outperform `eSS` in terms of speed and computational efficiency while keeping its robustness. Our design philosophy aligns closely with the principles of recently proposed “matheuristics”, i.e. combining rigorous mathematical frameworks with heuristic methods to create a more robust and efficient strategy for solving complex optimization problems [86].

## 3. Methods

### 3.1. Mathematical Formulation

Differential equations are fundamental tools for describing and predicting the evolution of variables over time. In systems biology, time-varying biological processes are typically modeled using systems of differential equations, collectively referred to as dynamic systems. Depending on the nature of the differential operators involved, these equations can be classified into several types, such as ordinary differential equations (ODEs), partial differential equations (PDEs), and differential-algebraic equations (DAEs).

In many biological applications, spatial effects are negligible or can be reasonably ignored. As a result, ODE-based models are commonly employed, as they capture the temporal dynamics of the system without incorporating spatial dimensions. Dynamic systems can also be classified based on the presence or absence of randomness. In deterministic dynamic systems, the system’s behavior is fully determined by the initial conditions and parameters, with no influence from random effects. In contrast, stochastic systems incorporate probabilistic elements to account for intrinsic or extrinsic noise.

Finally, a dynamic system is said to be mechanistic when it is grounded in physical, chemical, or biological laws that explain the underlying processes driving system behavior. Here, we consider the case of deterministic, mechanistic dynamic systems described by a system of ODEs:

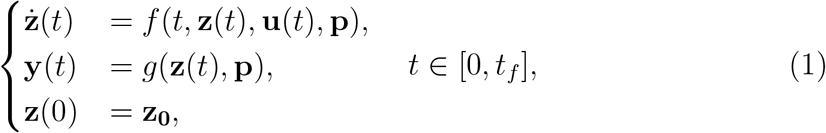

where:

- **z** is an *n*-dimensional vector representing the state variables, for some natural number *n*, and **z**_0_ describes its initial value.
- 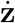 describes the time-derivative function of **z**.
- **u** is an *n*_*u*_-dimensional vector representing the time-varying input (control) variables, for some natural number *n*_*u*_.
- **y** is an *n*_*obs*_-dimensional vector representing the observable variables, for some natural number n_*obs*_ ≤ *n*.
- **p** is an m-dimensional vector representing the model parameters, for some natural number *m*.
- t denotes the time variable.
- 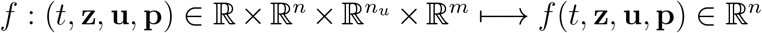 describes the system dynamics.
- 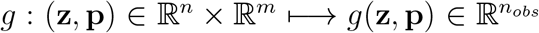, the observability function, describes the value of the observed variables.

In most cases, a dataset with measurements of the dynamic system is available, but the corresponding values of the model’s parameter vector remain unknown. Estimating these parameters is essential to preserve the model’s predictive capability. To achieve this, one typically defines an objective function that quantifies the discrepancy between the experimental data and the model’s predictions for a given set of parameters. The goal is then to identify the parameter values that minimize this function, subject to the dynamic system’s equations. This constitutes the well-known parameter estimation problem.

Two primary statistical frameworks can be used to define an appropriate objective function: the frequentist and the Bayesian approaches. In the frequentist view, the objective function typically quantifies the likelihood of the observed data given a particular parameter set. Conversely, the Bayesian approach incorporates prior beliefs or knowledge about the parameters’ distributions, introducing a probabilistic prior term into the formulation [2]. In most practical cases, there is limited or no prior information available regarding the distribution of the unknown parameters. Therefore, we adopt a frequentist perspective in this work.

Lat us now consider the formulation of the parameter estimation problem as a global optimization, where **x** ∈ ℝ^*m*^ is the vector of decision variables corresponding to the model parameters **p**. A commonly used objective function under this framework is the weighted least squares criterion, defined as follows:

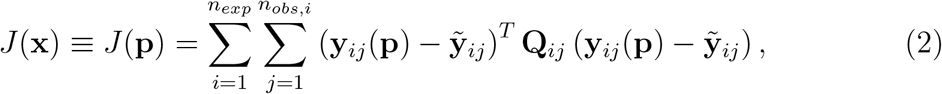

where:

- *n*_*exp*_ is the number of experiments,
- *n*_*obs,i*_ is the number of measured observables in experiment *i*.
- 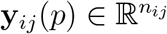 is the model prediction for observable *j* in experiment *i*,
- 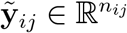 denotes the corresponding measured data,
- 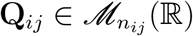 is a symmetric, non-negative definite weighting matrix.

This formulation allows the inclusion of multiple experiments and observables while assigning different levels of importance to each dataset, depending on the weighting matrices. It is important to note that the search domain for the parameter vector is typically not the entire space ℝ^*m*^. First and foremost, any candidate parameter set must satisfy the underlying dynamic system, as defined by Equation 1. Moreover, it is common for each parameter to be restricted to lie within predefined lower and upper bounds. In such cases, the problem is referred to as a bound-constrained dynamic optimization problem, which is the type of problem addressed in this work.

The evaluation of *J*(**p**) requires repeated numerical integration of the ODE system, typically via a single-shooting approach [14, 19, 60]. This strategy requires the solution of the inner initial value problem (the dynamic system described by ODEs) for each evaluation of the objective function. As a consequence, the resulting optimization problem is computationally expensive, highly nonlinear, multimodal, and often very ill-conditioned [9, 11]. These characteristics make parameter estimation in systems biology particularly challenging and motivate the use of robust global optimization methods.

### 3.2. The Proposed Algorithm: eLSHADE+

#### 3.2.1. Rationale and Baseline Method

As reviewed in previous sections, a broad range of global optimization methods has been proposed for parameter estimation in dynamic biological models. Among them, population-based metaheuristics have gained particular prominence due to their robustness, scalability, and limited reliance on gradient information, which may be unavailable or unreliable in this context.

As an initial step, we screened several state-of-the-art metaheuristics that have demonstrated strong performance in recent large-scale optimization studies and international benchmarking competitions. The selection criteria included scalability with problem dimension, robustness across heterogeneous landscapes, limited need for problem-specific tuning, and efficiency under strict computational budgets.

This screening indicated that methods based in Differential Evolution (`DE`) were particularly competitive. Among them, the Success-History based Adaptive Differential Evolution with Linear Population Size Reduction (`LSHADE`) [87] consistently ranked among the top-performing algorithms. `LSHADE` is an adaptive variant of Differential Evolution that extends the original `SHADE` algorithm by introducing a linear population size reduction mechanism. Its key innovation lies in the use of historical memories to adapt the crossover rate (*CR*) and scaling factor (*F*) based on parameter values that have led to successful offspring in previous iterations.

The algorithm employs a *current-to-*p*-best* mutation strategy combined with binomial crossover and a standard DE selection rule. An external archive stores replaced individuals, which may later be reused during mutation to promote diversity and mitigate premature convergence. As the optimization progresses, the population size is reduced linearly, gradually shifting the balance from exploration toward intensification.

However, preliminary experiments on parameter estimation problems from systems biology revealed that `LSHADE` did not achieve statistically significant improvements over the enhanced Scatter Search (`eSS`) method [20], which is widely regarded as the current state of the art in this domain. While `LSHADE` effectively addresses high-dimensional problems, its intensification mechanisms remain implicit and primarily driven by population contraction, which may be insufficient to efficiently refine solutions in the complex, rugged landscapes typical of biological models.

#### 3.2.2. New mechanisms

To address these limitations, we developed a novel hybrid approach, `eLSHADE+`, designed to surpass the performance of `eSS` while preserving the robustness and scalability of `LSHADE` (see contribution 1 in Section 1.1). The guiding principles were: (i) explicit control of the exploration–exploitation balance, (ii) sustained intensification throughout the optimization process, (iii) improved exploration of ill-scaled parameter spaces, and (iv) minimal increase in algorithmic complexity and user-defined hyperparameters.

##### Probabilistic Multi-Strategy Hybridization

The efficacy of hybrid metaheuristics is critically dependent on the synergy between global exploration and local exploitation. However, in the context of ODE-based parameter estimation, the objective function landscape exhibits a pathological duality. The underlying kinetic laws imply smooth, differentiable valleys where gradient-based methods converge rapidly. Conversely, numerical noise from ODE solvers (especially in stiff regimes) and structural non-identifiability create artificial roughness and multimodal traps where gradient methods fail [11]. Furthermore, biological landscapes are topologically heterogeneous, containing regions where any local search is computationally wasteful or prone to stagnation.

Standard hybrid metaheuristics commit to a single local solver, often failing when the landscape topology shifts between smooth and rugged regions. To address this fundamental limitation, `eLSHADE+` implements a probabilistic selection mechanism that probabilistically chooses among three operational modes at each local search trigger:

###### 1. Gradient-based Intensification

Employs `fmincon` from the MATLAB Optimization Toolbox [88], specifically using the interior-point algorithm. This mode exploits the differentiable structure of kinetic laws to achieve high-precision convergence in smooth basins of attraction, providing superior speed and accuracy when the landscape is locally differentiable.

###### 2. Derivative-free Robust Search

Uses Dynamic Hill Climbing (`dhc`) [89], a zero-order algorithm designed to handle rough, non-smooth, or noisy objective function landscapes. This mode provides robustness against numerical artifacts and discontinuities that break gradient solvers, ensuring reliability when the optimization trajectory traverses irregular regions of the parameter space.

###### 3. Pure Global Exploration (Null Strategy)

Deliberately skips the local search phase entirely. This mode preserves computational budget for the global evolutionary search, which is advantageous in complicated or fractal-like regions where local basins are narrow and deceptive. By avoiding premature intensification in suboptimal regions, this strategy prevents the waste of expensive function evaluations on refinement that yields marginal improvement.

The selection of these specific local solvers was guided by extensive prior benchmarking in the context of enhanced Scatter Search (`eSS`) [12, 20, 22–24, 61], which consistently identified `dhc` and `fmincon` as particularly competitive in terms of robustness, computational efficiency, and accuracy for systems biology problems.

Unlike approaches where local optimization is confined to final stages, our method performs local searches intermittently throughout the optimization, maintaining consistent intensification. The probabilistic switching strategy allows the algorithm to capture the complementary strengths of different local optimization paradigms while mitigating their individual limitations, particularly critical given the expensive nature of ODE-based objective function evaluations.

##### Logarithmic transformation of the search space

Parameter estimation problems in systems biology often involve parameters spanning several orders of magnitude (e.g., reaction rates from 10^−6^ to 10^3^). In such cases, exploration in the original parameter space is inefficient, as standard sampling strategies allocate equal density across the logarithmic range. Following established practices [12, 31, 59, 60, 90], we incorporate an optional logarithmic change of variables that transforms the search to log-space, facilitating uniform exploration across scales. This transformation can significantly accelerate convergence when parameter bounds differ by multiple orders of magnitude and can be enabled or disabled by the user based on prior knowledge of the problem structure.

#### 3.2.3. Algorithmic Workflow

The proposed method, termed `eLSHADE+`, is a hybrid global optimization algorithm that extends `LSHADE` by incorporating periodic local search and optional exploration in logarithmic space. Its overall workflow is summarized in Algorithm 1 and illustrated in Figure 1, while the local search procedure is detailed in Algorithm 2.

**Figure 1.**
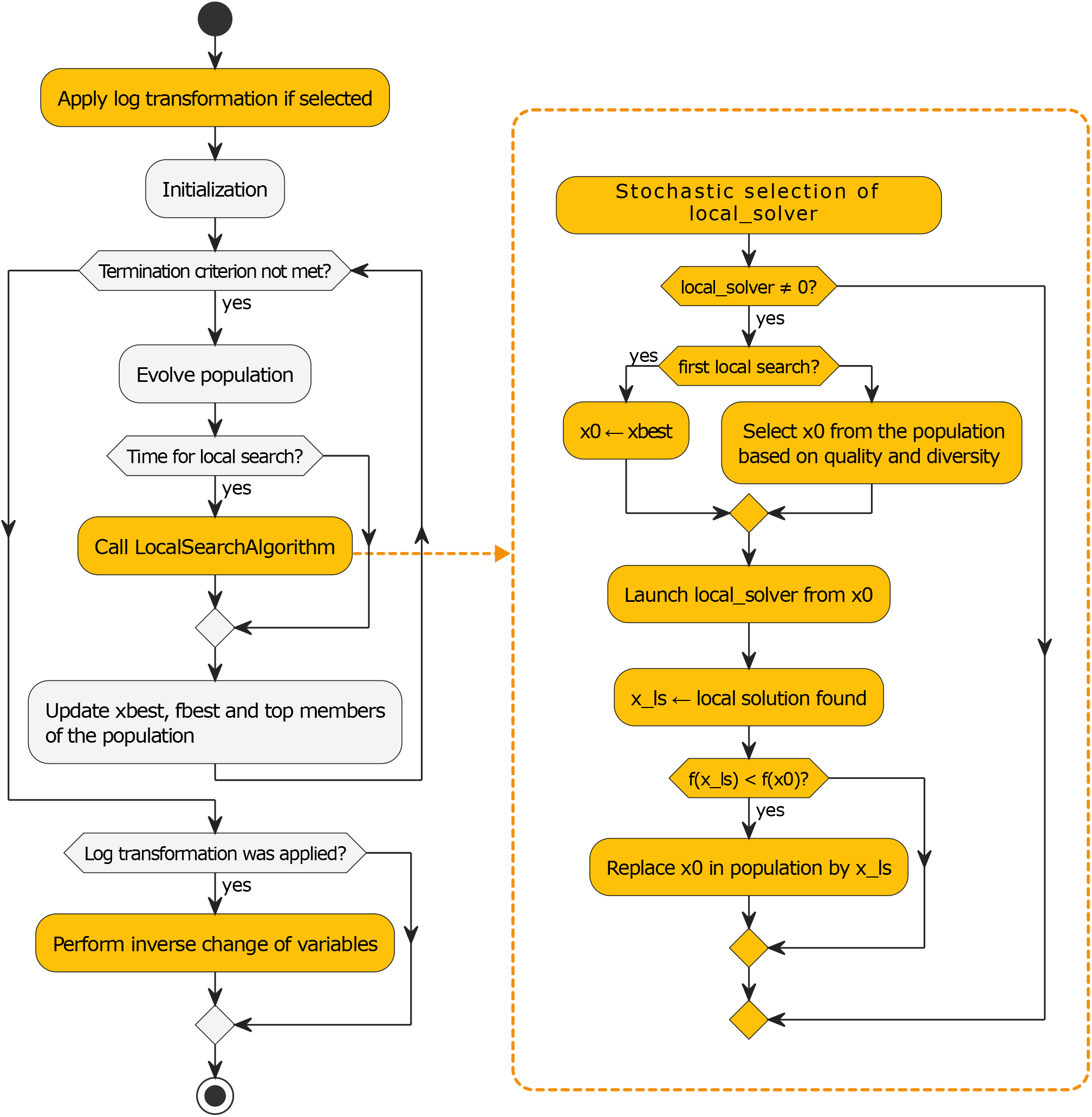
Simplified workflow of `eLSHADE+`.

At the beginning of the optimization process, the algorithm performs a logarithmic change of variables in case this option was enabled by the user. Next, an initial population *P* of *N*_*P,max*_ candidate solutions is generated within the feasible domain, and all individuals are evaluated. The best solution found so far, ***x***_*best*_, is stored, as well as its objective function value, *J*_*best*_. An external archive *A* is initialized as empty, and two historical memories of size *H* are initialized to generate crossover rates and scaling factors (*M*_*CR*_, *M*_*F*_). Additionally, an empty set *L* is created to store the initial points and local optima identified during the local search phase.

The algorithm then enters its main iterative loop. At each iteration, two temporary sets are initialized to store the crossover rates and scaling factors associated with successful offspring (*S*_*CR*_, *S*_*F*_). For each individual ***x***_*i*_ in the population, a crossover rate *CR*_*i*_ is sampled from a Gaussian distribution centered at a random element of *M*_*CR*_, while a scaling factor *F*_*i*_ is sampled from a Cauchy distribution centered at a random element of *M*_*F*_.

Mutation is performed using the *current-to-*p*-best* strategy. Let ***x***_*pbest*_ be one of the *p* · *N*_*P*_ individuals in the population, based on objective function values. The mutant vector is generated by the following formula:

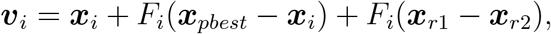

where ***x***_*r*1_ is a random member of P, and ***x***_*r*2_ is a random member of *P* ∪*A*. Binomial crossover is then applied to generate a trial vector ***u***_*i*_, ensuring that at least one component of the mutant vector is inherited. If ***u***_*i*_ yields a lower objective function value than ***x***_*i*_, it replaces ***x***_*i*_ in the population. In this case, ***x***_*i*_ enters *A*, and *CR*_*i*_ and *F*_*i*_ are stored in *S*_*CR*_ and *S*_*F*_, respectively.

Then, A is pruned if its size exceeds the predefined maximum. If at least one successful offspring has been generated during the iteration, *M*_*CR*_ and *M*_*F*_ are updated using the weighted Lehmer mean of the values stored in *S*_*CR*_ and *S*_*F*_, following the standard `LSHADE` mechanism. The population is truncated according to the linear reduction of its size.

Every *n*_*ls*_ iterations, the algorithm triggers a local search phase. The optimization strategy is selected probabilistically: a derivative-free method is selected with probability *w*_1_, and a gradient-based method is selected with probability *w*_2_. The local search is skipped with probability 1 − (*w*_1_ + *w*_2_). The starting point, ***x***_*start*_, for the local search is selected based on a rank that takes into consideration both quality and diversity, balancing intensification around high-quality solutions with exploration of previously unvisited regions. The trade-off between intensification and diversification is determined by a balance parameter that ranges from 0 to 1. If the balance parameter is set to 0, the rank is based solely on the objective function values of the individuals (measurement of quality). If set to 1, the rank relies only on the distance of individuals to *L* (measurement of diversity). The initial point ***x***_*start*_ and the local solution found, ***x***_*ls*_, are added to L. If ***x***_*ls*_ improves the objective function value of ***x***_*start*_, it replaces it in *P*. At the end of each iteration, the top *p* · *N*_*P*_ members of *P* are updated.

The iterative process continues until a predefined termination criterion is met. If a logarithmic transformation was applied at the beginning of the optimization, the inverse transformation is performed to recover the solution in the original parameter space.

### 3.3. Ablation study and configuration selection

To quantify the contribution of individual components within the proposed `eLSHADE+` framework and to justify the selection of its default configuration, we conducted a systematic ablation study. This analysis isolates the impact of three primary design choices: (i) the use of a logarithmic transformation of the parameter space, (ii) the inclusion of a periodic local search mechanism, and (iii) the specific selection strategy for the local solver. The workflow for these components corresponds to Algorithms 1 and 2.

#### Algorithm 1

`eLSHADE+`

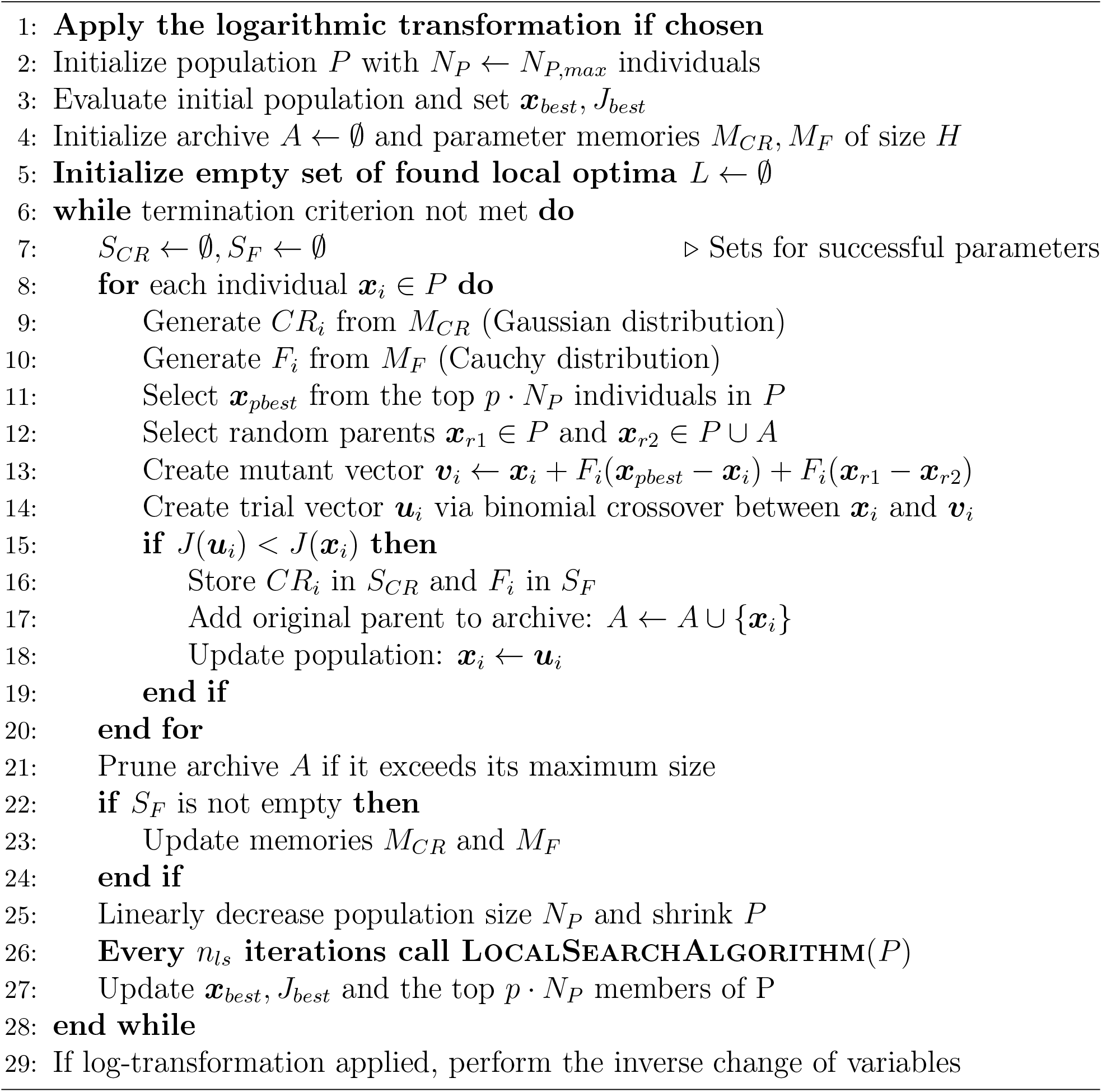

#### Algorithm 2

LocalSearchAlgorithm

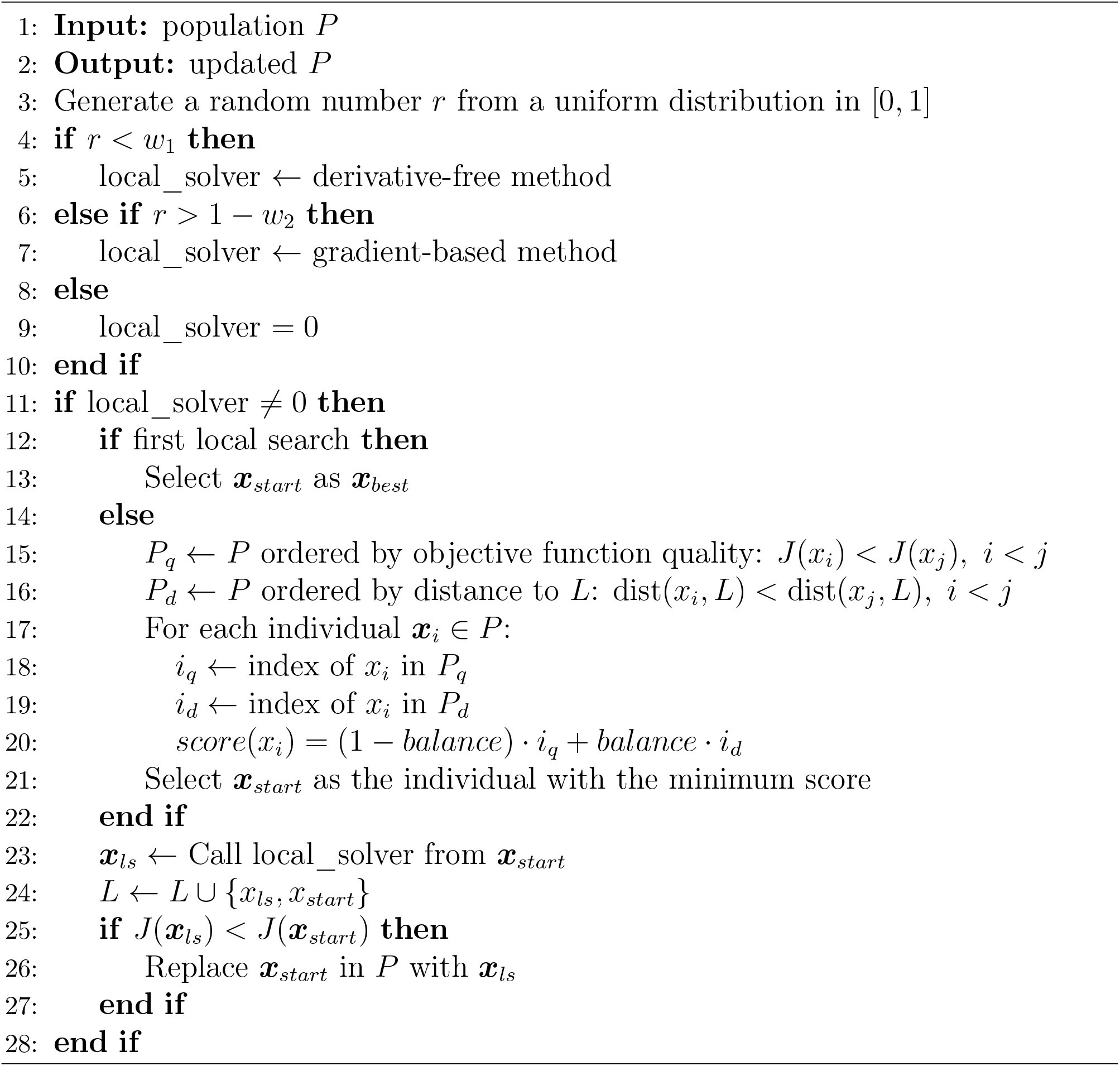

While the logarithmic transformation is controlled by a binary toggle, the local search configuration involves multiple parameters. The algorithm triggers a local search phase every *n*_*ls*_ iterations. At this interval, a probabilistic decision determines whether to execute a local refinement and, if so, which strategy to employ. We consider two solvers with distinct information requirements:

1. A derivative-free method: Dynamic Hill Climbing (`dhc`) [89].
2. A gradient-based method: The `fmincon` algorithm from MATLAB [88].

The selection is governed by two weights, *w*_1_ and *w*_2_. In every local search cycle, a random number *r* ∼ *U*[0, 1] is generated. The derivative-free method (`dhc`) is selected if *r* ∈ [0, *w*_1_], while the gradient-based method (`fmincon`) is selected if *r* ∈ [1 − *w*_2_, 1]. If neither condition is met, the local search phase is skipped. Therefore, varying w_1_ and w_2_ allows us to modulate the hybridization intensity and the balance between gradient-based and derivative-free local refinement.

Table 1 details the eight configurations evaluated. The `Ablation Baseline` represents the proposed default setup, where logarithmic sampling is active and the algorithm maintains an equiprobable balance between exploring via `dhc`, refining via `fmincon`, or continuing with global search (i.e., *w*_1_ = *w*_2_ = 1/3). Alternative configurations systematically isolate these factors by strictly enforcing a specific local solver, disabling local search entirely, or deactivating the logarithmic transformation.

**Table 1:** Different configurations of the proposed hybrid method considered on the Ablation Study. For each configuration, we specify whether the logarithmic search is applied, as well as the values of *w*_1_ and *w*_2_ (weights for the selection of the method to be applied in the local search phase).

| Ablation Study: Tested Configurations |  |  |
| --- | --- | --- |
| Configuration | Logarithmic Search | Local Search |
| A. Ablation Baseline | Activated | Activated: $w_1 = 1/3, w_2 = 1/3$. |
| B. Log + dhc | Activated | Activated: $w_1 = 1, w_2 = 0$. |
| C. Log + fmincon | Activated | Activated: $w_1 = 0, w_2 = 1$. |
| D. Log + NoLS | Activated | Deactivated: $w_1 = 0, w_2 = 0$. |
| E. NoLog + EqWeights | Deactivated | Activated: $w_1 = 1/3, w_2 = 1/3$. |
| F. NoLog + dhc | Deactivated | Activated: $w_1 = 1, w_2 = 0$. |
| G. NoLog + fmincon | Deactivated | Activated: $w_1 = 0, w_2 = 1$. |
| H. NoLog + NoLS | Deactivated | Deactivated: $w_1 = 0, w_2 = 0$. |

### 3.4. Benchmark Problems

The assessment of global optimization algorithms is often performed using synthetic benchmark functions. While such functions are useful for controlled comparisons, they fail to capture many of the structural and numerical challenges encountered in real-world optimization problems, including complex constraint interactions, ill-conditioning, heterogeneous parameter sensitivities, and expensive objective function evaluations. These limitations are particularly pronounced in parameter estimation for dynamic biological models, where the objective function arises from the numerical solution of nonlinear differential equations [22, 82].

For this reason, we evaluate the proposed method using the BioPreDyn benchmark set [62], a collection of large-scale, biologically motivated parameter estimation problems specifically designed to reflect the complexity of realistic dynamic systems. The benchmark includes models of varying size, topology, and experimental design, and has been widely adopted in the literature as a reference for assessing global optimization methods in systems biology.

An important characteristic of the BioPreDyn benchmark is that its primary objective is to evaluate the ability of optimization algorithms to minimize the objective function, rather than to ensure accurate recovery of parameter values. This design choice reflects the fact that large-scale biological models often exhibit severe structural and practical identifiability issues, making unique parameter estimation infeasible even under ideal conditions. Consequently, algorithmic performance is assessed based on objective function values and convergence behavior, rather than parameter identifiability.

Table 2 summarizes the main characteristics of the five benchmark problems included in the BioPreDyn set. Together, these problems span a wide range of dimensionalities, from medium-scale models with fewer than 100 parameters to a genome-scale system with over 1,700 parameters.

**Table 2:** Main features of the BioPreDyn Benchmark set [62].

| Problem | B1 | B2 | B3 | B4 | B5 |
| --- | --- | --- | --- | --- | --- |
| **Organism** | *S. cerevisiae* | *E. coli* | *E. coli* | CHO cells | Generic |
| **Number of parameters** | 1759 | 116 | 178 | 117 | 86 |
| **Dynamic states** | 276 | 18 | 47 | 34 | 26 |
| **Observed states** | 44 | 9 | 47 | 13 | 6 |

#### Problem B1

This benchmark represents a genome-scale metabolic network of *Saccharomyces cerevisiae*. The reaction kinetics are described using a generalized form of reversible Michaelis–Menten equations. The model comprises 261 reactions, 262 dynamic variables, and 1,759 parameters to be estimated. Experimental data consist of 44 steady-state measurements, which were verified to be stable. However, these data were insufficient for reliable parameter estimation. To address this limitation, synthetic data were generated using a Monte Carlo sampling procedure within predefined parameter bounds. For each observable, 120 samples were generated and contaminated with uniform noise at a level of 5%.

#### Problem B2

This model describes the central carbon metabolism of *Escherichia coli* and its dynamic response to a glucose pulse. The system includes 18 state variables, 48 reactions, and 116 unknown parameters. Calibration was performed using experimental data obtained from real measurements, making this problem representative of realistic experimental conditions.

#### Problem B3

This benchmark models the adaptation of *Escherichia coli* to changes in carbon sources across three consecutive growth phases: glucose, acetate, and a mixture of both. The model consists of 47 ordinary differential equations, with all state variables being observable. A total of 178 parameters are estimated. Calibration was performed using pseudo-experimental data without added noise, allowing the evaluation of algorithmic performance under idealized conditions.

#### Problem B4

This model represents the metabolism of Chinese Hamster Ovary (CHO) cells, which are widely used in biotechnological fermentation processes. The system includes 35 metabolites, 32 reactions, and 117 parameters. Pseudo-experimental data were generated by adding Gaussian noise with a standard deviation of 5% to simulated observations.

#### Problem B5

The final benchmark corresponds to a signal transduction logic model composed of 26 ordinary differential equations. State variables represent protein activity levels and are constrained to lie between 0 and 1. A total of 86 parameters are estimated. Calibration data were generated in silico across 10 experiments, each measuring 6 observables. For each observable, 16 equally spaced time points were sampled, and Gaussian noise with variable standard deviation was added to simulate experimental measurement error.

#### Additional problems B1_e, B2_e, and B4_e

Additionally, we considered modified versions of benchmarks B1, B2, and B4, for which the upper bounds were set to be 100 times larger than their original values. These new versions are more difficult to solve, as the search domain is wider. We denote these problems as B1_e, B2_e, and B4_e, respectively. Considering these variants is useful for demonstrating the effectiveness of logarithmic transformations when the problem involves wide bounds, which is often the case in real-world scenarios where there is no prior knowledge of the parameter ranges and the context requires exploring extensive search domains.

### 3.5. Experimental Setup and Comparative Framework

To evaluate the performance of the proposed `eLSHADE+` algorithm, we compared it against a selection of established global optimization methods commonly applied to parameter estimation in systems biology [13, 19, 22, 24, 31, 55, 81]. These methods were chosen based on their proven effectiveness in handling multimodal, highdimensional, and ill-conditioned problems. They span several algorithmic families:

- **Deterministic Global Optimization:** We include `DIRECT` [91], which provides a baseline for systematic exploration of the search space without randomized heuristics. This contrasts with the stochastic nature of the remaining algorithms.
- **Clustering and Multi-start Methods:** Algorithms such as `MLSL` [92] employ clustering techniques to identify basins of attraction, effectively combining global sampling with local refinement while avoiding redundant searches in previously explored regions.
- **Randomized Search:** `CRS` [93, 94] is a classical randomized search algorithm that has shown good performance in certain problem classes.
- **Evolutionary and Population-based Strategies:** The majority of the benchmarked methods rely on population-based stochastic search. These include:
  - *Genetic and Differential Algorithms:* Standard `GA` [95] and the adaptive Differential Evolution variant `LSHADE` [87, 96, 97], which mimic biological evolution through recombination and mutation.
  - *Evolution Strategies (ES):* `CMA-ES` [98], `ISRES` [99], and `ESCH` [100, 101], which generally focus on self-adapting mutation distributions (such as covariance matrices) to navigate ill-conditioned landscapes.
  - *Enhanced Scatter Search:* `eSS` [20] can be interpreted as a population-based method that utilizes an advanced scatter search framework designed to systematically combine solutions, acting as a bridge between pure evolutionary exploration and local exploitation.

This heterogeneity ensures that the proposed method is tested against solvers with different balances of exploration (diversification) and exploitation (intensification). More details about these methods and their implementation are available at [25].

#### 3.5.1. Implementation and Computational Environment

Numerical integration of the BioPreDyn ODE models for all methods was performed using the CVODES solver from the SUNDIALS suite [102] to ensure consistent objective function evaluation and used C-compiled MEX files to ensure computational efficiency [62]. In order to manage the optimizations with the different methods using a common interface, we developed `MEIGO2-alfa`, a MATLAB toolbox based on `MEIGO` [103] that provides a unified framework and facilitates the benchmarking. Code and comprehensive documentation, including implementation dependencies and usage instructions, are provided in our Zenodo repository [25]. Table 3 summarizes the primary references and source implementations for the algorithms integrated within the `MEIGO2` environment

**Table 3:**
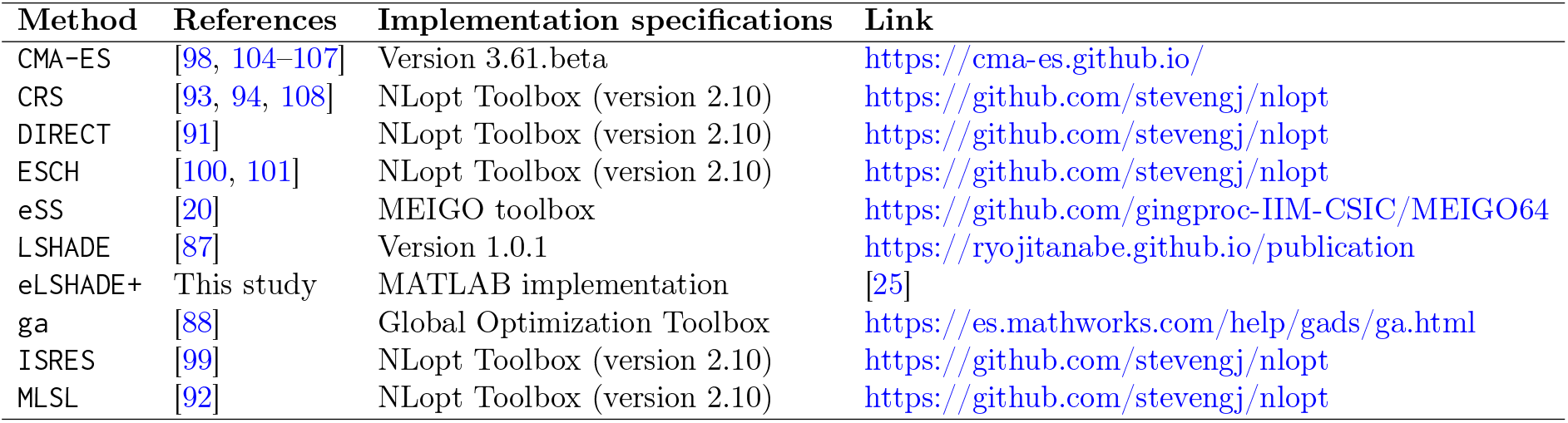
Overview of optimization methods compared in this study, including key references and implementation sources used for their integration on MEIGO2-alfa.

The computational experiments for benchmarks B1, B2, B4, and B5 were conducted on a workstation equipped with an AMD EPYC 8434P 48-Core Processor (96 logical threads, 2.5 GHz) and 128 GB of RAM, running Windows 11 Pro. Benchmark B3, due to specific restrictions regarding the encoding of the objective function, was executed on the DRAGO Linux cluster hosted at CSIC, utilizing Intel Xeon Gold 6248R processors. According to Passmark CPU benchmarks (single thread), the AMD and Intel CPUs have similar performance. The MATLAB version employed in both infrastructures was R2024b (Update 5).

#### 3.5.2. Algorithm Configuration and Experimental Protocol

Prior to the main study, a preliminary screening was conducted on standard mathematical optimization functions to tune the proposed method. For the core evaluation on the five biological benchmarks, each problem was solved in 10 independent runs to account for stochastic variability.

Parameter settings were adopted from literature recommendations, with the exception of benchmark B1, where the population size was adjusted for `eSS`, `eLSHADE+` and `LSHADE` to accommodate its high dimensionality. Table 4 lists the key hyperparameters for the evolutionary algorithms (note that `DIRECT` is excluded as the NLopt implementation does not permit customization). To prevent premature termination and ensure a fair assessment of convergence capability, all stopping criteria other than the maximum execution time were disabled.

**Table 4:**
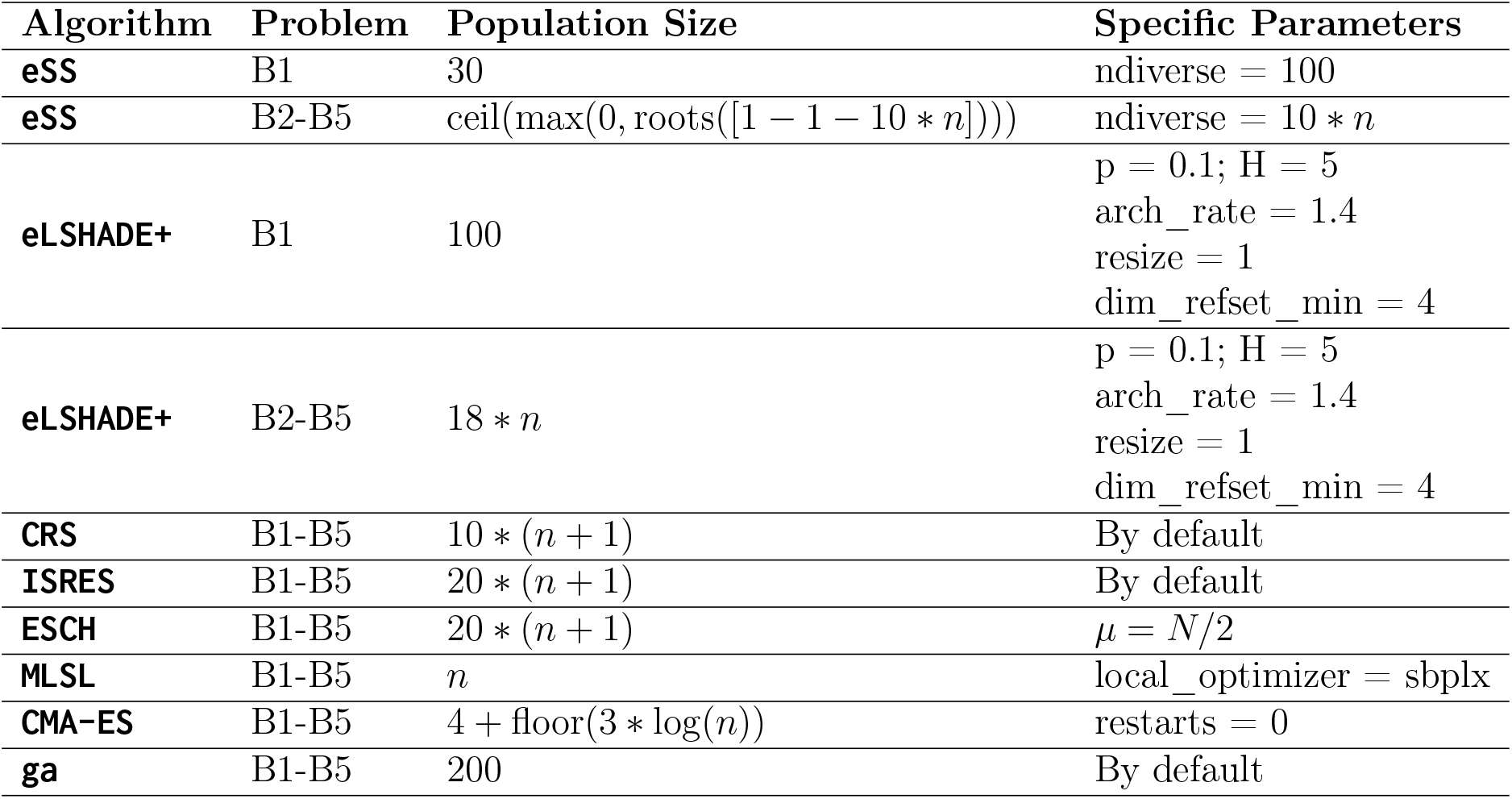
Parameter settings for the algorithms considered. The configuration of LSHADE coincides with that for eLSHADE+. *n* denotes the number of decision variables (parameters) in the model.

| Algorithm | Problem | Population Size | Specific Parameters |
| --- | --- | --- | --- |
| eSS | B1 | 30 | ndiverse = 100 |
| eSS | B2-B5 | $\text{ceil}(\max(0, \text{roots}([1 - 1 - 10 * n])))$ | ndiverse = $10 * n$ |
| eLSHADE+ | B1 | 100 | p = 0.1; H = 5<br>arch_rate = 1.4<br>resize = 1<br>dim_refset_min = 4 |
| eLSHADE+ | B2-B5 | $18 * n$ | p = 0.1; H = 5<br>arch_rate = 1.4<br>resize = 1<br>dim_refset_min = 4 |
| CRS | B1-B5 | $10 * (n + 1)$ | By default |
| ISRES | B1-B5 | $20 * (n + 1)$ | By default |
| ESCH | B1-B5 | $20 * (n + 1)$ | $\mu = N/2$ |
| MLSL | B1-B5 | $n$ | local_optimizer = sbplx |
| CMA-ES | B1-B5 | $4 + \text{floor}(3 * \log(n))$ | restarts = 0 |
| ga | B1-B5 | 200 | By default |

#### 3.5.3. Metrics for Comparison

Comparing algorithm performance in this domain requires standardized criteria. We primarily analyze the *convergence curves*, which plot the best-so-far objective function value against computation time. This allows for a “vertical cut” analysis at the maximum time threshold, identifying which algorithm achieved the lowest error within the budget. This time-based approach is preferred over counting function evaluations, as it accounts for the overhead of algorithmic logic and implementation differences, reflecting real-world utility.

Additionally, we use boxplots to visualize the dispersion of the final objective values, offering insight into the robustness of the methods. Statistical analysis is provided via the arithmetic mean (sensitive to outliers/failures) and the median (representative of typical performance), along with the standard deviation to quantify variability. To confirm statistical significance, we applied the Wilcoxon rank-sum test to pairwise comparisons of the top five methods for each benchmark.

To quantify performance beyond raw objective function values, we define two additional normalized metrics. Let 𝒜 be the set of algorithms, and *P* the problem. For an algorithm *A* of 𝒜, we denote by *J*_*A,P*_ the set of objective function values achieved in each run of algorithm *A* for problem *P*, and by med(*J*_*A,P*_) the median of the aforementioned set. The following metric *m*_1_ accounts for normalized performance relative to the best and worst algorithms considered:

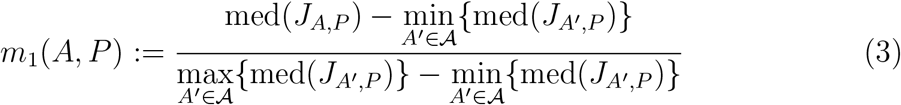

The second metric, *m*_2_, measures the logarithmic deviation from the nominal optimal value for problem *P* (where available), which we denote by *J*_*nom,P*_ :

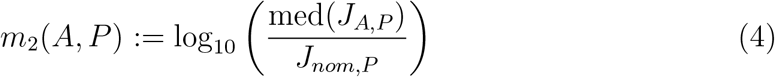

Lower values indicate better performance for both metrics. For visualization, *m*_2_ values were linearly mapped to the unit interval [0, 1].

#### 3.5.4. Data and Code Availability

All source code, benchmark implementations, experimental results, and supplementary materials are publicly available in a Zenodo repository [25]. The repository includes:

- Complete implementation of `eLSHADE+` in MATLAB
- An integrated software suite with all comparative algorithms
- Data, code interface and results for all benchmarks (B1–B5 and extended variants). Statistical analysis with p-value tables are also included
- Ablation study results
- Detailed documentation and usage instructions

DOI: https://doi.org/10.5281/zenodo.18379327

## 4. Results

This section presents the numerical results obtained across the considered benchmark problems. We first provide the results of the ablation study. In the following subsection, we provide a summary of the aggregate results across all benchmarks. Next, we conduct a detailed problem-specific analysis, focusing on the five top-performing algorithms for each case. Comprehensive results, including raw data and additional visualizations, are available in the Supplementary Material and our online repository [25].

### 4.1. Results of the ablation study

Experimental validation was performed using the BioPreDyn benchmark suite (problems B1–B5) [62]. To increase the complexity of the test landscape, we also considered the modified versions of problems B1, B2 and B4 with extended parameter bounds described in subsection 3.4 and denoted by B1_e, B2_e, and B4_e, respectively. Therefore, this ablation study was performed over a total of eight optimization problems. Each configuration underwent 10 independent runs per problem.

Table 5 summarizes the results obtained through this study. They confirm that the `Ablation Baseline` systematically yields the best performance overall, not only with respect to the reduction of the objective function but also in terms of robustness, presenting the lowest standard deviations among all tested configurations. The results obtained for B1_e, B2_e, and B4_e confirm that exploration in logarithmic space is highly recommendable when parameter bounds are wide. This motivates selecting the ablation baseline as the default configuration of our algorithm. Detailed numerical data is provided in the supplementary file `Ablation_Study_eLSHADE+.xlsx` available in our repository [25].

**Table 5:** Summary of results of the ablation study. Configuration labels correspond to those detailed in Table 1 Values within 5% of the lowest median for each problem are shown in bold.

|  |  | Configuration |  |  |  |  |  |  |  |
| --- | --- | --- | --- | --- | --- | --- | --- | --- | --- |
|  |  | A | B | C | D | E | F | G | H |
| B1 | Median | **8.3420E+03** | 9.3622E+03 | 9.6902E+03 | **8.2719E+03** | 9.6030E+03 | 1.1042E+04 | 1.0330E+04 | **8.0844E+03** |
|  | std | 6.5398E+02 | 1.4578E+03 | 7.7235E+02 | 6.8972E+02 | 2.2132E+03 | 1.7513E+04 | 1.3697E+04 | 1.6893E+03 |
| B1_e | Median | 1.5960E+04 | 1.4860E+04 | 1.4860E+04 | **9.6540E+03** | 3.4349E+04 | 3.7689E+04 | 3.5502E+04 | 1.9177E+04 |
|  | std | 3.4976E+03 | 5.3213E+03 | 5.3213E+03 | 2.0157E+03 | 7.8863E+03 | 1.0724E+04 | 8.4315E+03 | 7.7764E+03 |
| B2 | Median | **2.2092E+02** | **2.2374E+02** | 2.3029E+02 | 2.2885E+02 | **2.2027E+02** | **2.1510E+02** | 2.4031E+02 | 2.3991E+02 |
|  | std | 1.2216E+01 | 1.3949E+01 | 1.2981E+01 | 2.1216E+01 | 2.0778E+01 | 1.7518E+01 | 1.4984E+01 | 1.8647E+01 |
| B2_e | Median | **1.3512E+02** | 2.0444E+02 | **1.3605E+02** | **1.3748E+02** | 2.0030E+02 | 1.4364E+02 | 5.0618E+02 | 7.8662E+02 |
|  | std | 5.0933E+01 | 6.4866E+01 | 5.1409E+01 | 3.8851E+01 | 3.2565E+02 | 3.4019E+02 | 3.3305E+02 | 2.9960E+02 |
| B3 | Median | **1.5444E-02** | 1.6906E-02 | 1.9055E-02 | 2.1342E-02 | 4.9802E-02 | 4.8540E-02 | 4.4696E-02 | 5.5541E-02 |
|  | std | 9.0985E-03 | 1.9355E-02 | 1.5164E-02 | 1.0468E-02 | 1.3398E-02 | 4.4036E-02 | 2.6550E-02 | 6.4247E+00 |
| B4 | Median | **3.3033E+01** | 3.3507E+01 | **3.3053E+01** | **3.2493E+01** | **3.1794E+01** | **3.2569E+01** | **3.2010E+01** | **3.2206E+01** |
|  | std | 7.1493E-01 | 1.0546E+00 | 5.4105E-01 | 4.4686E-01 | 8.7949E-01 | 9.8553E-01 | 3.9128E-01 | 8.5162E-01 |
| B4_e | Median | 1.0738E+02 | 7.5842E+01 | 5.3239E+01 | **4.7212E+01** | 1.8871E+05 | 3.4416E+04 | 7.9506E+05 | 6.1337E+03 |
|  | std | 4.3206E+03 | 3.5404E+03 | 2.2660E+03 | 1.1267E+02 | 7.1113E+05 | 8.4895E+04 | 1.3200E+06 | 1.0751E+06 |
| B5 | Median | **2.8820E+03** | **2.8787E+03** | **2.8797E+03** | **2.8821E+03** | **2.9112E+03** | **2.9412E+03** | 3.0369E+03 | 3.0419E+03 |
|  | std | 2.6827E+02 | 2.6738E+02 | 2.0359E+02 | 3.3480E+02 | 5.2676E+01 | 8.1695E+01 | 5.5154E+02 | 3.0231E+02 |

### 4.2. Overall Comparison of Methods

Table 6 summarizes the primary outcomes for the BioPreDyn benchmarks. It identifies the top-performing algorithm, the computational time budget, the nominal (optimal) objective value *J*_*nom*_ (where known), the best value across all runs of the top-performing algorithm *J*_*best*_, and the best value achieved in previous literature (*J*_*ref*_). Notably, the results obtained in this study consistently surpass the reference values from [62].

**Table 6:** Summary of results for each benchmark. For each, we specify: the top-performing algorithm (achieving the lowest objective function value across all runs); the computational time budget (used as the stopping criterion); *J*_*nom*_: the nominal value for the objective function corresponding to the nominal parameters; *J*_*best*_: the objective function value reached by the top-performing algorithm in this study; *J*_*ref*_ : the best value reported by [62].

| Model | B1 | B2 | B3 | B4 | B5 |
| --- | --- | --- | --- | --- | --- |
| Algorithm | LSHADE | eLSHADE+ | eLSHADE+ | LSHADE | eLSHADE+ |
| Time budget (h) | 24 | 6 | 48 | 6 | 20 |
| $J_{nom}$ | 1.2564E+06 | - | 0 | 3.9071E+01 | 4.2727E+03 |
| $J_{best}$ | **7.6011E+03** | **1.9763E+02** | **7.1772E-03** | **3.1363E+01** | **2.8761E+03** |
| $J_{ref}$ | 1.3753E+04 | 2.3390E+02 | 3.7029E-01 | 4.5718E+01 | 3.0725E+03 |

**Table 7:** Summary of results for benchmark B1. For each algorithm, we report: the best value of all runs, and the mean, median and standard deviation of all the solutions reported. We highlight: the lowest best solution found, the lowest mean, and the lowest median.

|  | Best | Mean | Median | std |
| --- | --- | --- | --- | --- |
| CMA-ES | 1.2196E+06 | 3.8025E+10 | 7.9120E+07 | 1.1945E+11 |
| CRS | 7.0372E+04 | 8.1074E+04 | 8.0001E+04 | 7.7755E+03 |
| DIRECT | 4.2145E+04 | 4.2145E+04 | 4.2145E+04 | 0.0000E+00 |
| ESCH | 1.0461E+05 | 2.8950E+07 | 9.7074E+05 | 6.6137E+07 |
| eSS | 9.1420E+03 | 1.0310E+04 | 9.9859E+03 | 1.3040E+03 |
| eSS+dhc | 9.1757E+03 | 1.0360E+04 | 1.0187E+04 | 8.1751E+02 |
| eSS+fmincon | 8.7988E+03 | 1.0409E+04 | 9.9310E+03 | 1.4853E+03 |
| LSHADE | **7.6011E+03** | 8.8590E+03 | **8.0844E+03** | 1.6893E+03 |
| eLSHADE+ | 7.8213E+03 | **8.6358E+03** | 8.3420E+03 | 6.5398E+02 |
| ga | 1.0334E+04 | 2.2407E+04 | 1.8976E+04 | 9.9115E+03 |
| MLSL | 8.4496E+03 | 6.5601E+09 | 5.4718E+06 | 1.4455E+10 |
| ISRES | 8.9172E+04 | 2.8214E+08 | 3.7266E+05 | 8.8265E+08 |

**Table 8:** Summary of results for benchmark B2. For each algorithm, we report: the best value of all runs, and the mean, median and standard deviation of all the solutions reported. We highlight: the lowest best solution found, the lowest mean, and the lowest median.

|  | Best | Mean | Median | std |
| --- | --- | --- | --- | --- |
| CMA-ES | 6.9494E+02 | 3.5826E+04 | 2.2900E+03 | 6.0484E+04 |
| CRS | 2.6527E+02 | 7.1347E+02 | 6.1805E+02 | 3.9042E+02 |
| DIRECT | 1.0783E+03 | 1.0783E+03 | 1.0783E+03 | 0.0000E+00 |
| ESCH | 2.8895E+02 | 3.3353E+02 | 3.1819E+02 | 4.6650E+01 |
| eSS | 2.1581E+02 | 2.4247E+02 | 2.4452E+02 | 1.0129E+01 |
| eSS+dhc | 2.3278E+02 | 2.4239E+02 | 2.4379E+02 | 5.8705E+00 |
| eSS+fmincon | 2.3150E+02 | 2.4764E+02 | 2.4680E+02 | 8.5921E+00 |
| LSHADE | 1.9994E+02 | 2.2821E+02 | 2.3991E+02 | 1.8647E+01 |
| eLSHADE+ | **1.9763E+02** | **2.2294E+02** | **2.2092E+02** | 1.2216E+01 |
| ga | 2.9121E+02 | 5.6559E+02 | 5.2574E+02 | 2.1598E+02 |
| MLSL | 2.9449E+02 | 3.2903E+02 | 3.2110E+02 | 3.3780E+01 |
| ISRES | 2.6424E+02 | 2.8465E+02 | 2.7763E+02 | 1.8577E+01 |

**Table 9:**
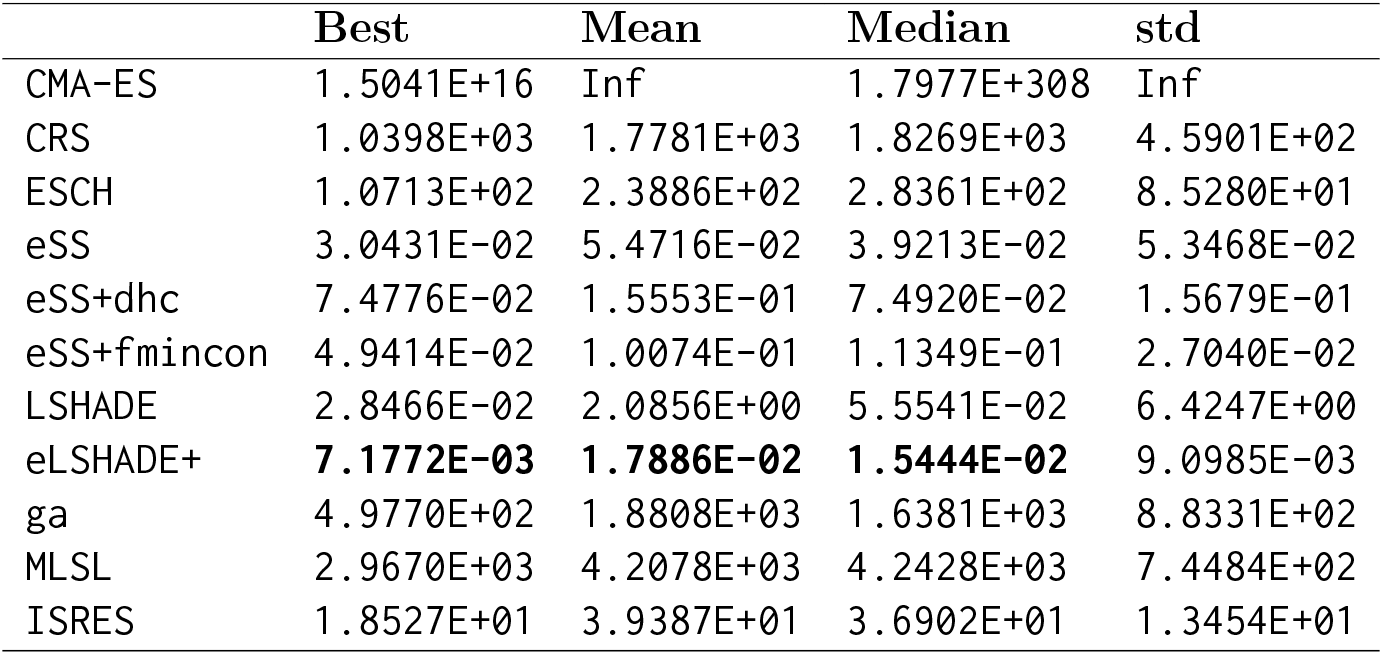
Summary of results for benchmark B3. For each algorithm, we report: the best value of all runs, and the mean, median and standard deviation of all the solutions reported. We highlight: the lowest best solution found, the lowest mean, and the lowest median.

|  | Best | Mean | Median | std |
| --- | --- | --- | --- | --- |
| CMA-ES | 1.5041E+16 | Inf | 1.7977E+308 | Inf |
| CRS | 1.0398E+03 | 1.7781E+03 | 1.8269E+03 | 4.5901E+02 |
| ESCH | 1.0713E+02 | 2.3886E+02 | 2.8361E+02 | 8.5280E+01 |
| eSS | 3.0431E-02 | 5.4716E-02 | 3.9213E-02 | 5.3468E-02 |
| eSS+dhc | 7.4776E-02 | 1.5553E-01 | 7.4920E-02 | 1.5679E-01 |
| eSS+fmincon | 4.9414E-02 | 1.0074E-01 | 1.1349E-01 | 2.7040E-02 |
| LSHADE | 2.8466E-02 | 2.0856E+00 | 5.5541E-02 | 6.4247E+00 |
| eLSHADE+ | **7.1772E-03** | **1.7886E-02** | **1.5444E-02** | 9.0985E-03 |
| ga | 4.9770E+02 | 1.8808E+03 | 1.6381E+03 | 8.8331E+02 |
| MLSL | 2.9670E+03 | 4.2078E+03 | 4.2428E+03 | 7.4484E+02 |
| ISRES | 1.8527E+01 | 3.9387E+01 | 3.6902E+01 | 1.3454E+01 |

Figure 2 illustrates (with boxplots) the distribution of final objective values. The left column displays all algorithms, while the right column focuses on the top five performers (lowest median values). The method presented here, `eLSHADE+`, was the best overall performer (in terms of the best median), although in several cases presented an spread which is not statiscally different from `LSHADE`. `eSS` was not the best method for any of these problems, but achieved the second best median in two of the problems, a reflection of its good performance for this class of problems. More detailed comparisons, including statistical tests, are reported below.

**Figure 2.**
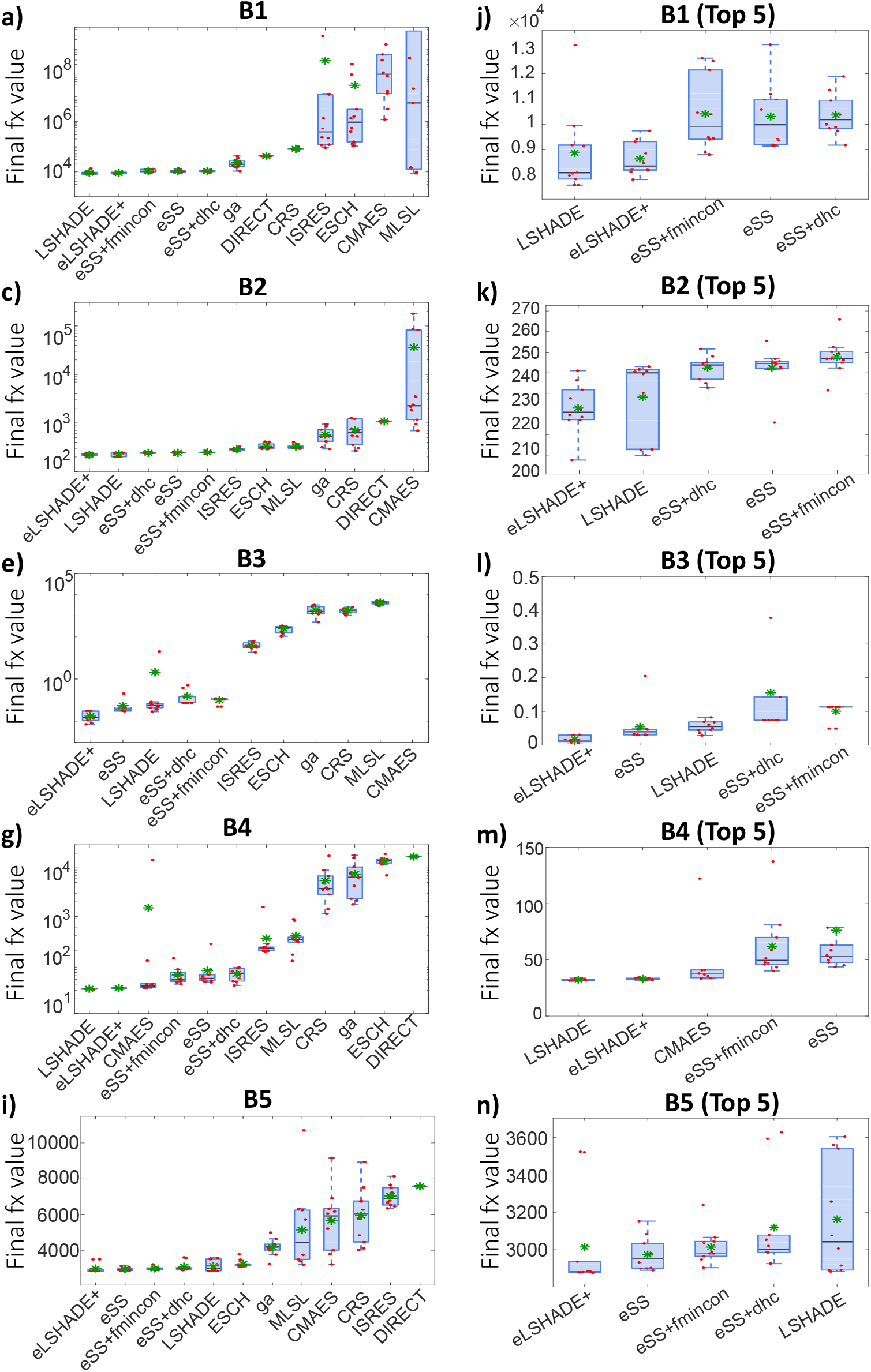
Boxplots of the final objective values for all algorithms on the BioPreDyn benchmark set. Algorithms are ordered left to right by increasing median.

Considering the additional normalized metrics, Figure 3 presents them via heatmaps and inverted spider plots for the best-performing methods (`LSHADE`, `eLSHADE+`, and `eSS` variants). These plots confirm that `eLSHADE+` consistently occupies the largest area in the spider plots, indicating superior overall performance across the benchmark set.

**Figure 3.**
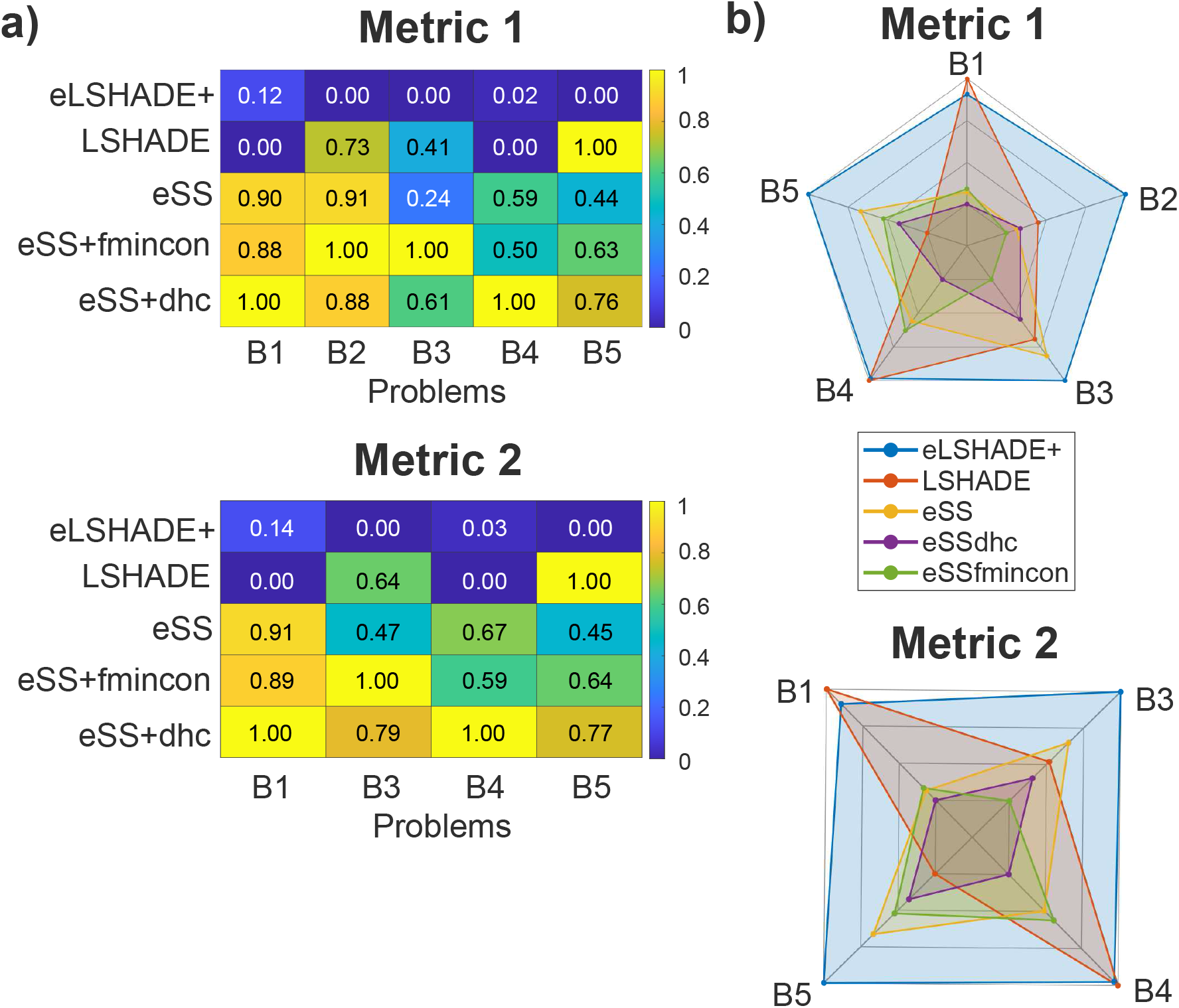
Heatmaps and spider plots for the computed metrics. In the spider plots, axes are inverted so that better performance corresponds to polygons with larger area. Figure a) illustrates the values of Metric 1 (normalized distance to the lowest median) and Metric 2 (discrepancy with respect to the nominal value). Figure b) corresponds to the spider plots of both metrics. Benchmark B2 is excluded for Metric 2 due to the absence of a nominal value.

### 4.3. Problem-Specific Comparison and Analysis

We now present a detailed analysis of the five algorithms that yielded the lowest median objective values for each benchmark. We utilize convergence plots to illustrate the optimization trajectory (both for the best individual run and the median across all runs) and report the outcomes of the Wilcoxon rank-sum tests to verify statistical significance (full *p*-values provided in our online repository [25]).

#### 4.3.1. Benchmark B1: Genome-scale Metabolic Network of S. cerevisiae

This problem represents a high-dimensional challenge (1,759 parameters). As shown in Figure 4, methods based on Differential Evolution (`DE`) significantly outperformed all other strategies. While our hybrid `eLSHADE+` achieved substantial error reduction and low dispersion, the original `LSHADE` algorithm yielded slightly superior results. The Wilcoxon test confirmed that both `LSHADE` and `eLSHADE+` were statistically superior to `eSS` and its hybrid variants, though the difference between the two DE variants was not statistically significant.

**Figure 4.**
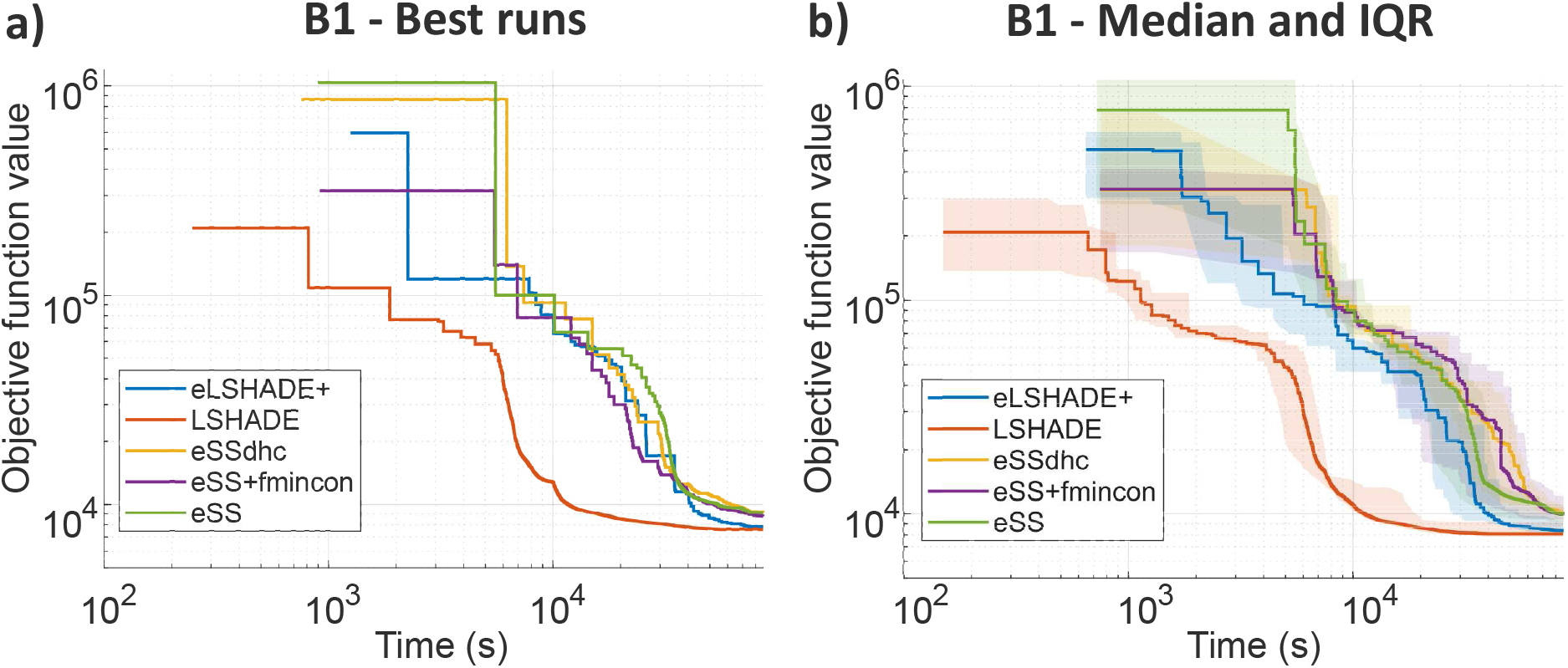
Evolution over time of the solutions reported by the top five algorithms for benchmark B1. Left: curves from the best run of each algorithm (i.e., the run with the lowest final-time solution). Right: curves showing the median and interquartile range across all runs.

A close inspection of the convergence curves reveals a slight advantage of the original `LSHADE` in this specific case, which is somewhat paradoxical. We believe this can be explained taking into account two effects which are particularly important in this problem: (i) in high-dimensional spaces where function evaluations are expensive, the overhead of the local search mechanism in `eLSHADE+` may reduce the time budget available for global exploration; and (ii) the logarithmic transformation is most effective in landscapes where the lower and upper bounds differ by several orders of magnitude, which is not the case for benchmark B1. Indeed, performing the logarithmic change of variables appears to introduce difficulties during the initial exploration phase for this specific problem, explaining the higher initial performance of the `LSHADE` curves. The results obtained by `eLSHADE+` and `LSHADE` on the modified problem B1_e confirm these hypotheses, demonstrating the superiority of `eLSHADE+` over `LSHADE` when the search domain is enlarged. These results are summarized in Table 12 and illustrated through Figures 5 and 6. This behavior validates our second contribution (Section 1.1): in high-dimensional spaces with expensive evaluations, selective intensification outperforms aggressive local refinement.

**Figure 5.**
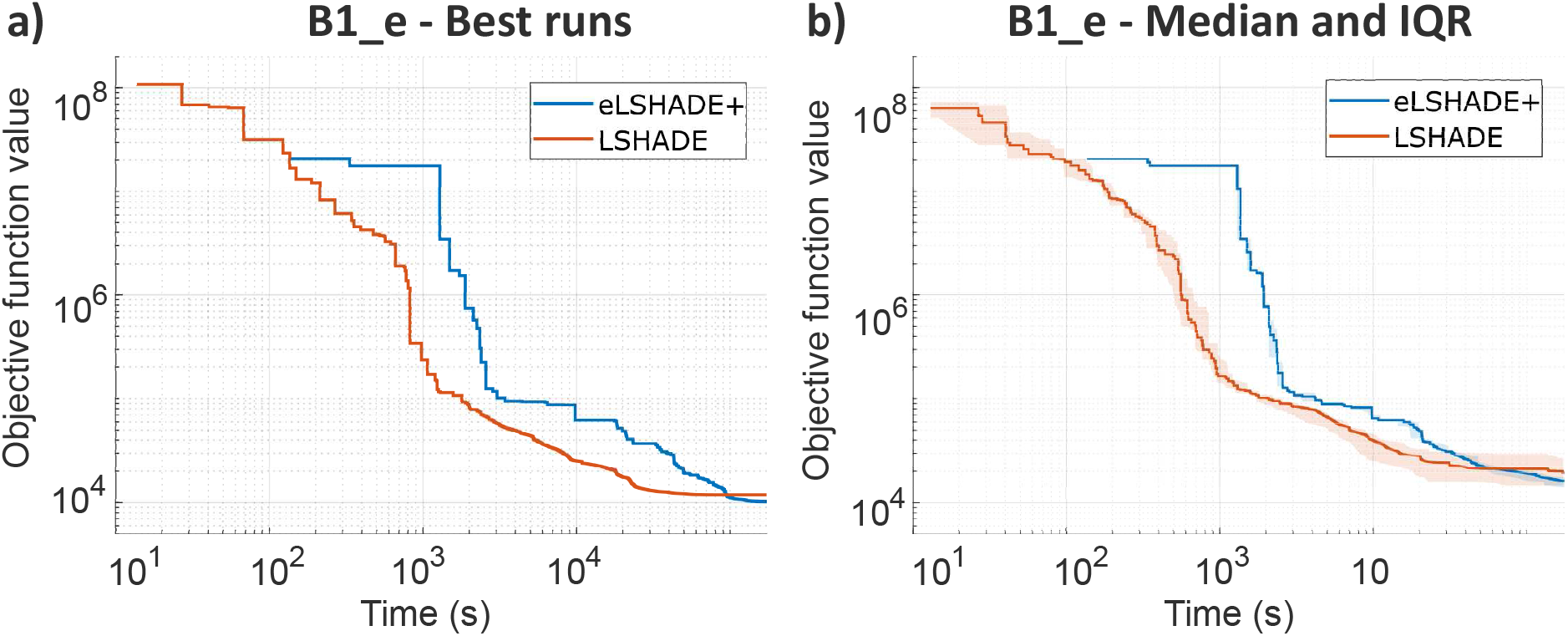
Evolution over time of the solutions reported by `eLSHADE+` and `LSHADE` for problem B1_e. Left: curves from the best run of each (i.e., the run with the lowest final-time solution). Right: curves showing the median and interquartile range across all runs.

**Figure 6.**
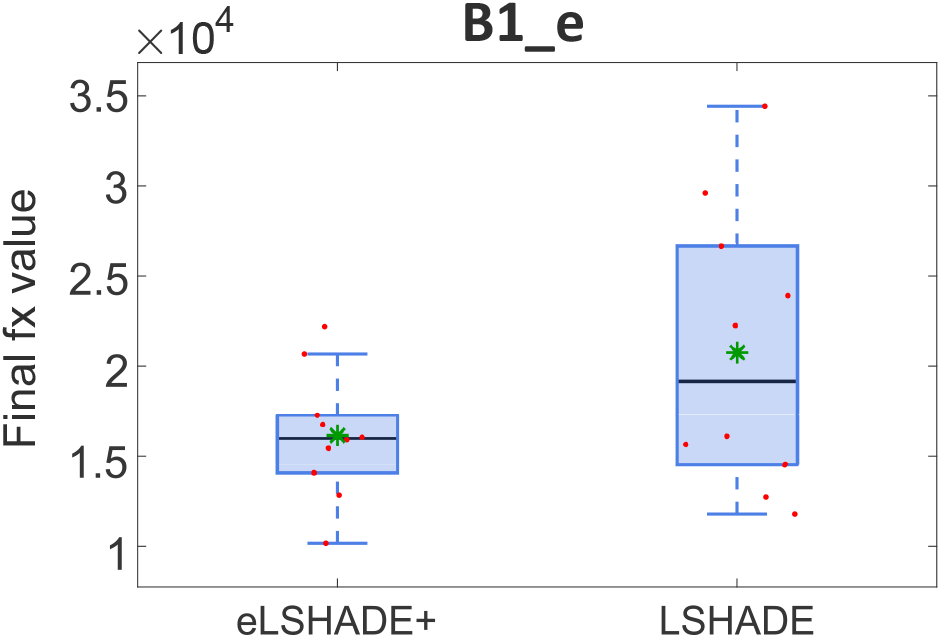
Boxplots of the final objective values for `eLSHADE+` and `LSHADE` for problem B1_e.

#### 4.3.2. Benchmark B2: E. coli Carbon Metabolism

For benchmark B2, `DE`-based methods again proved statistically superior. Figure 7 reveals an interesting dynamic: while `eSS` and its hybrid variants demonstrated faster initial convergence, `eLSHADE+` ultimately converged to the lowest objective function values (see also Figure 2d). This highlights the capability of `eLSHADE+` to escape local optima that trap scatter search methods in the later stages of optimization. Once again, the results for the modified version of B2, namely B2_e, highlight how the benefits of `eLSHADE+` are accentuated in wide search domain scenarios, as evidenced by Figures 8 and 9, and the results shown in Table 12.

**Figure 7.**
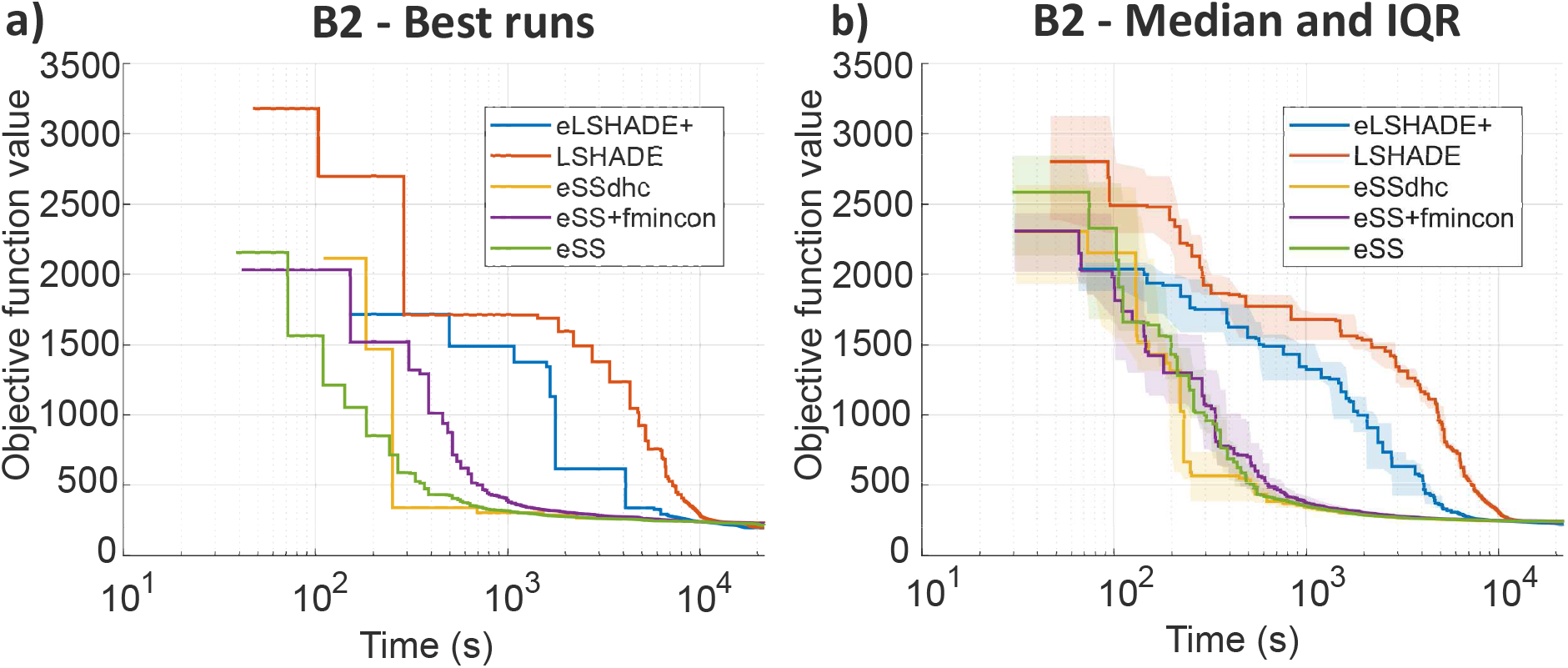
Evolution over time of the solutions reported by the top five algorithms for benchmark B2. Left: curves from the best run of each algorithm (i.e., the run with the lowest final-time solution). Right: curves showing the median and interquartile range across all runs.

**Figure 8.**
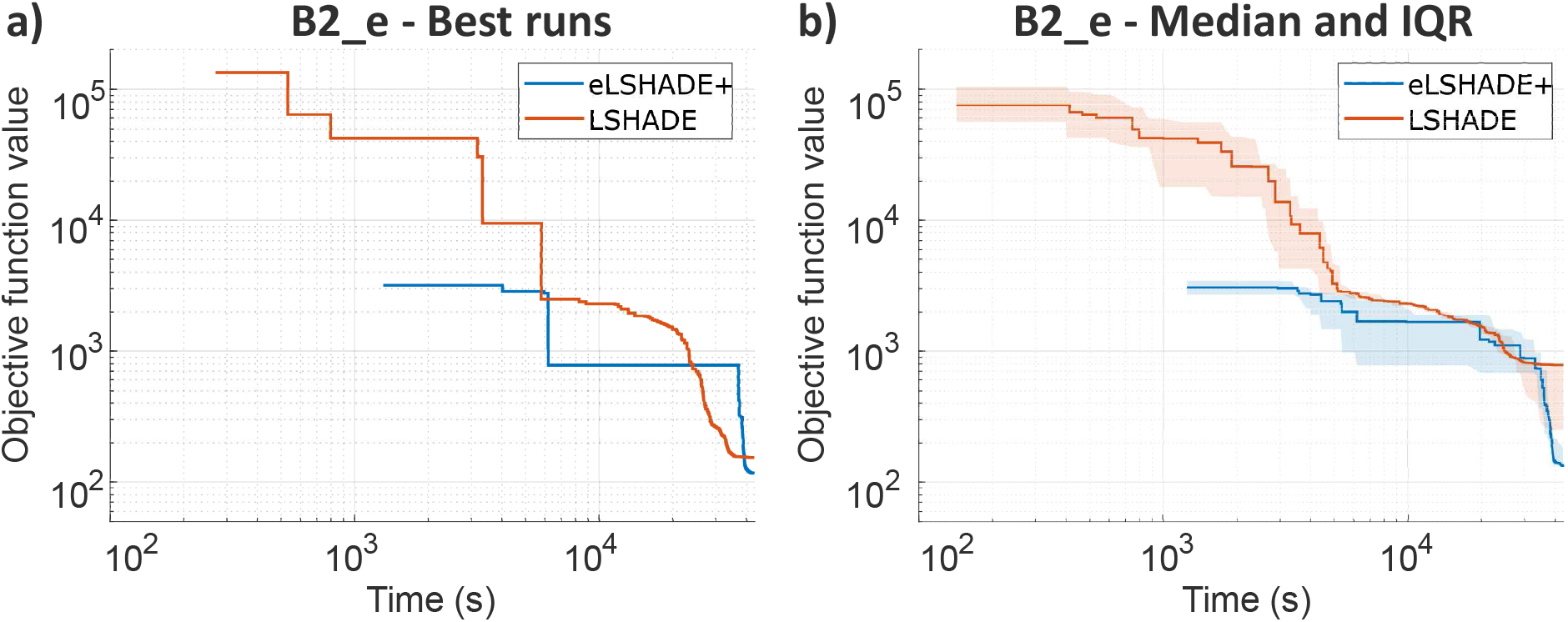
Evolution over time of the solutions reported by `eLSHADE+` and `LSHADE` for problem B2_e. Left: curves from the best run of each (i.e., the run with the lowest final-time solution). Right: curves showing the median and interquartile range across all runs.

**Figure 9.**
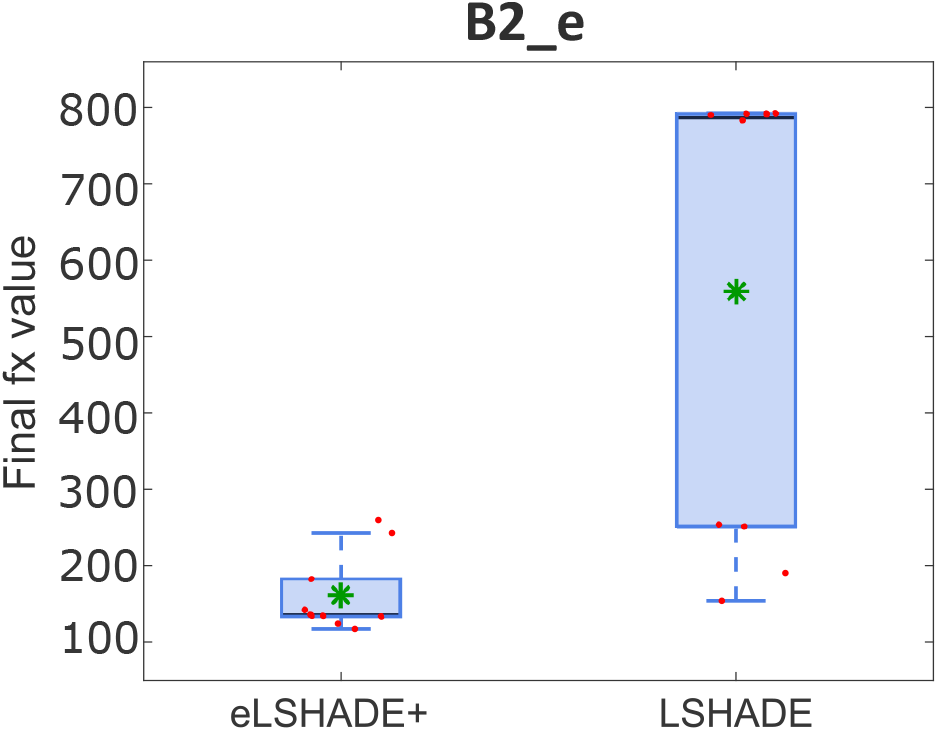
Boxplots of the final objective values for `eLSHADE+` and `LSHADE` for problem B2_e.

#### 4.3.3. Benchmark B3: E. coli Growth Adaptation

Benchmark B3 was the most computationally demanding problem (48 hours per run). Here, `eLSHADE+` exhibited a distinct advantage, returning the best values overall with very low dispersion (Figure 10 and Figure 2f). The efficacy of the logarithmic search space transformation was particularly pronounced here, as the parameter bounds spanned significantly different orders of magnitude. The Wilcoxon test confirmed that `eLSHADE+` performed statistically better than all other compared methods.

**Figure 10.**
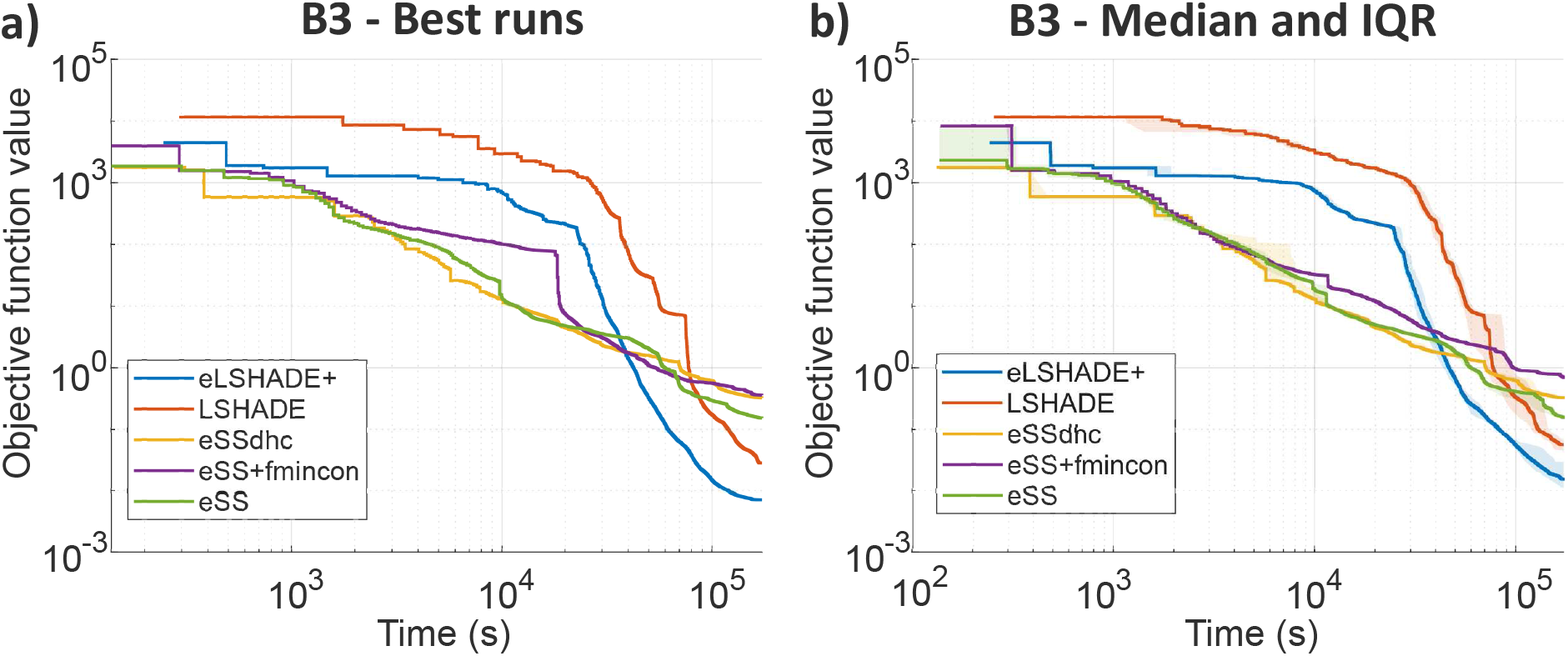
Evolution over time of the solutions reported by the top five algorithms for benchmark B3. Left: curves from the best run of each algorithm (i.e., the run with the lowest final-time solution). Right: curves showing the median and interquartile range across all runs.

#### 4.3.4. Benchmark B4: CHO Metabolism

In this benchmark, `CMA-ES` appeared in the top five. Note that the apparent delay in the starting time for `eLSHADE+`, `LSHADE`, and `eSS` in Figure 11 is an artifact of reporting protocols: these population-based methods only report a solution after evaluating the entire initial population, whereas `CMA-ES` reports from the first evaluation. Despite this, `LSHADE` and `eLSHADE+` achieved the lowest objective values and demonstrated superior robustness (lowest standard deviation, Table 10), outperforming `eSS` and its variants.

**Table 10:** Summary of results for benchmark B4. For each algorithm, we report: the best value of all runs, and the mean, median and standard deviation of all the solutions reported. We highlight: the lowest best solution found, the lowest mean, and the lowest median.

|  | Best | Mean | Median | std |
| --- | --- | --- | --- | --- |
| CMA-ES | 3.3224E+01 | 1.5014E+03 | 3.7507E+01 | 4.6018E+03 |
| CRS | 1.1285E+03 | 5.5010E+03 | 3.8284E+03 | 4.8965E+03 |
| DIRECT | 1.7126E+04 | 1.7126E+04 | 1.7126E+04 | 0.0000E+00 |
| ESCH | 7.0066E+03 | 1.3810E+04 | 1.4131E+04 | 3.1479E+03 |
| eSS | 4.3716E+01 | 7.6286E+01 | 5.3122E+01 | 6.8623E+01 |
| eSS+dhc | 3.7814E+01 | 6.5426E+01 | 6.7738E+01 | 1.8707E+01 |
| eSS+fmincon | 4.0167E+01 | 6.2356E+01 | 4.9813E+01 | 2.9325E+01 |
| LSHADE | **3.1363E+01** | **3.2318E+01** | **3.2206E+01** | 8.5160E-01 |
| eLSHADE+ | 3.2092E+01 | 3.3130E+01 | 3.3033E+01 | 7.1490E-01 |
| ga | 1.7766E+03 | 7.6311E+03 | 6.4353E+03 | 5.7767E+03 |
| MLSL | 1.2085E+02 | 3.9887E+02 | 3.3579E+02 | 2.5100E+02 |
| ISRES | 1.8972E+02 | 3.5329E+02 | 2.2518E+02 | 4.2734E+02 |

**Figure 11.**
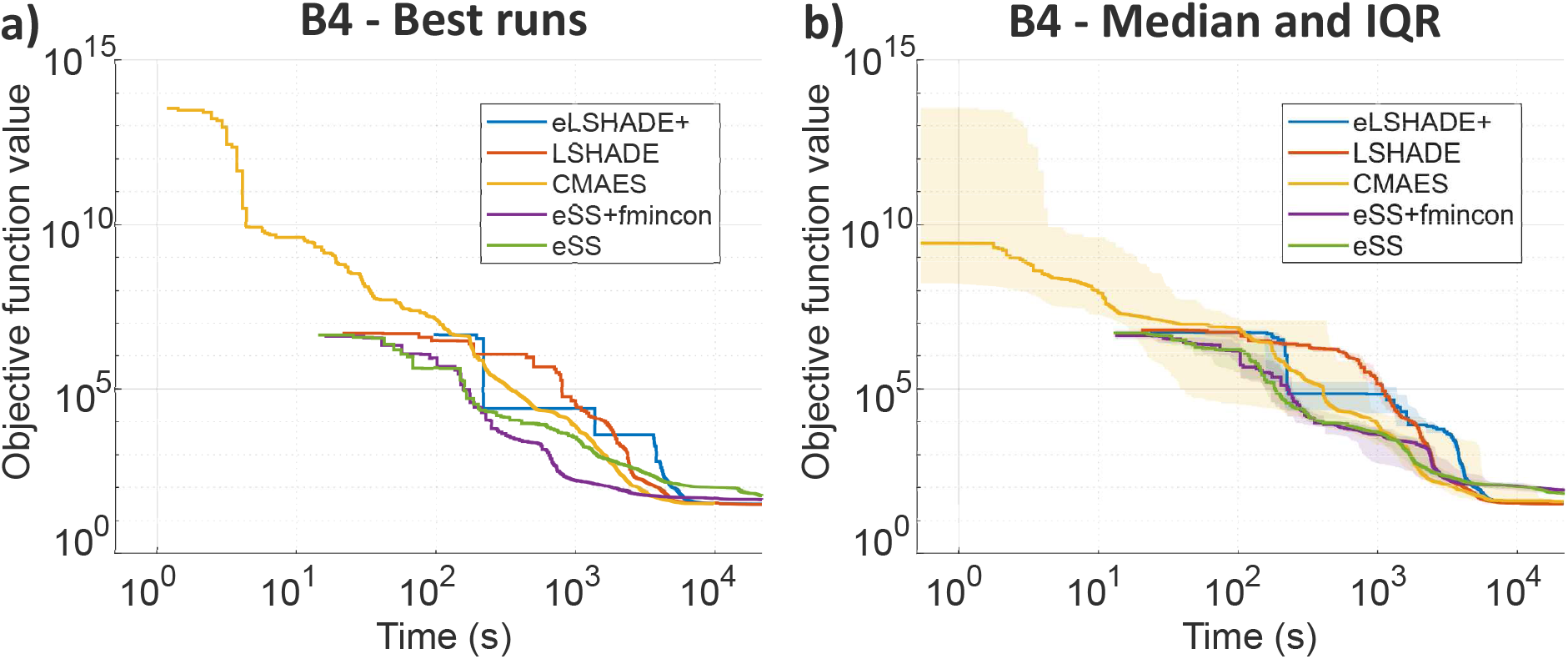
Evolution over time of the solutions reported by the top five algorithms for benchmark B4. Left: curves from the best run of each algorithm (i.e., the run with the lowest final-time solution). Right: curves showing the median and interquartile range across all runs.

Although the solutions reported by `eLSHADE+` and `LSHADE` appear similar, their performance differs significantly when the modification with extended bounds, B4_e, is considered, as shown in Figures 12 and 13, and Table 12.

**Figure 12.**
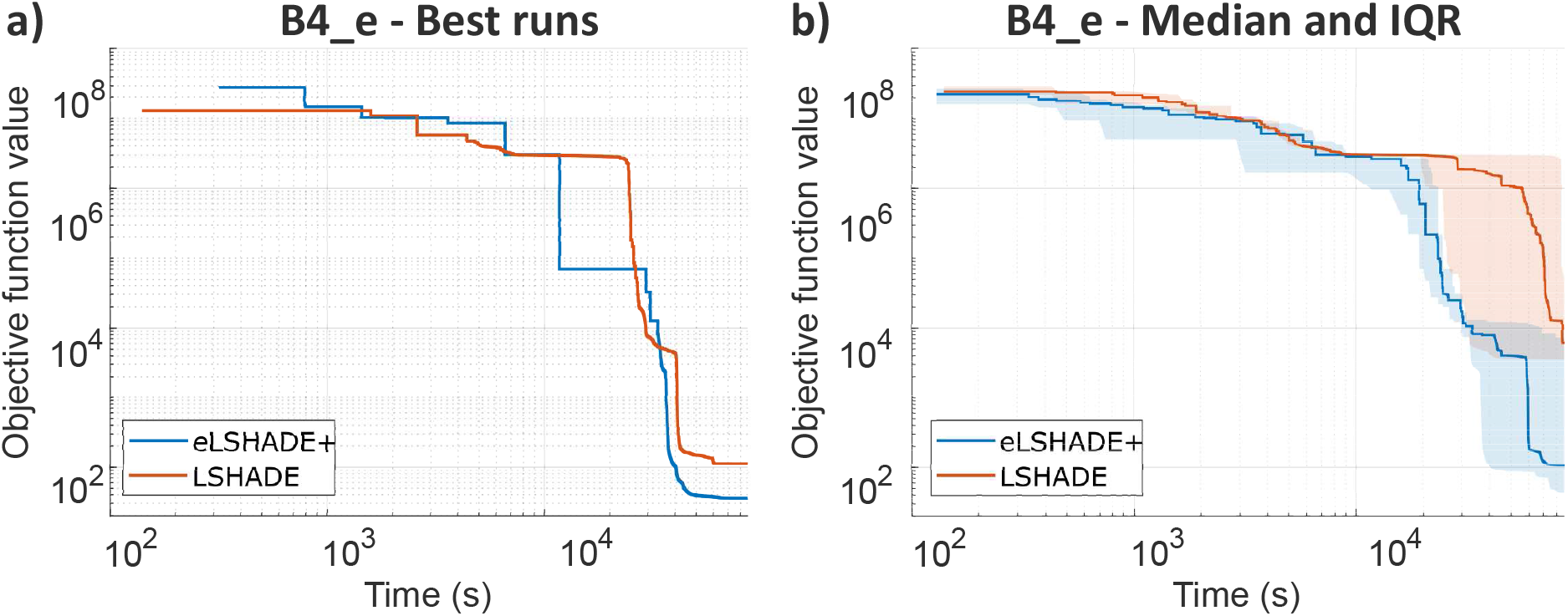
Evolution over time of the solutions reported by `eLSHADE+` and `LSHADE` for problem B4_e. Left: curves from the best run of each (i.e., the run with the lowest final-time solution). Right: curves showing the median and interquartile range across all runs.

**Figure 13.**
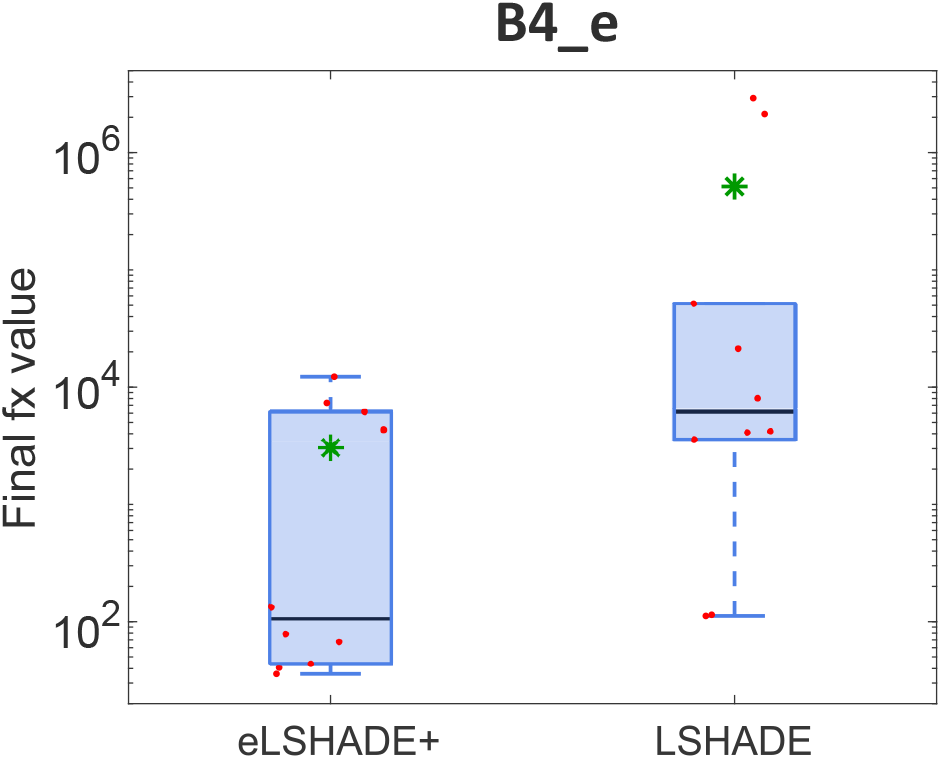
Boxplots of the final objective values for `eLSHADE+` and `LSHADE` for problem B4_e.

**Figure 14.**
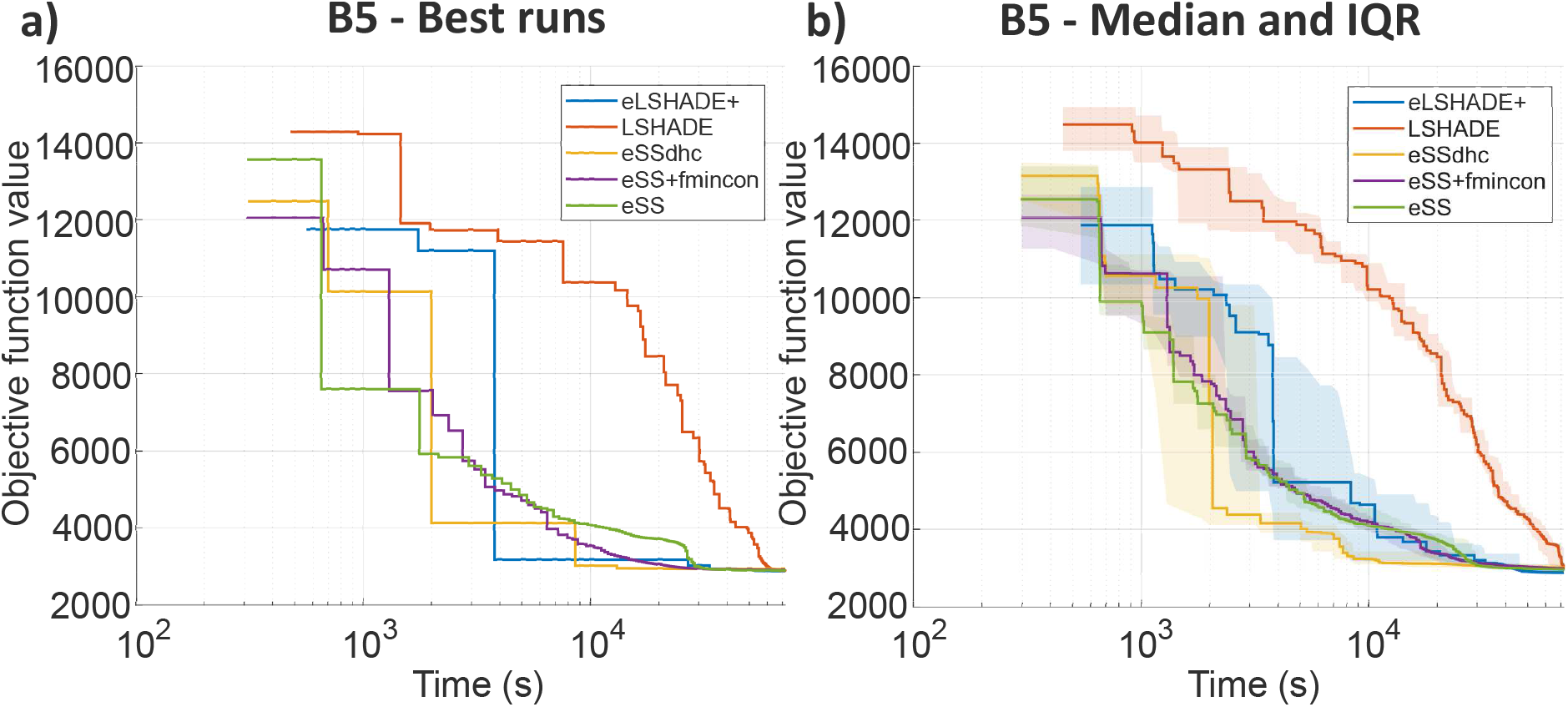
Evolution over time of the solutions reported by the top five algorithms for benchmark B5. Left: curves from the best run of each algorithm (i.e., the run with the lowest final-time solution). Right: curves showing the median and interquartile range across all runs.

#### 4.3.5. Benchmark B5: Signal Transduction Logic

Finally, for benchmark B5, `eLSHADE+` was the only method to produce statistically superior results compared to the rest of the set, as confirmed by the Wilcoxon test. Table 11 highlights its robustness, maintaining a low standard deviation where other methods fluctuated significantly.

**Table 11:**
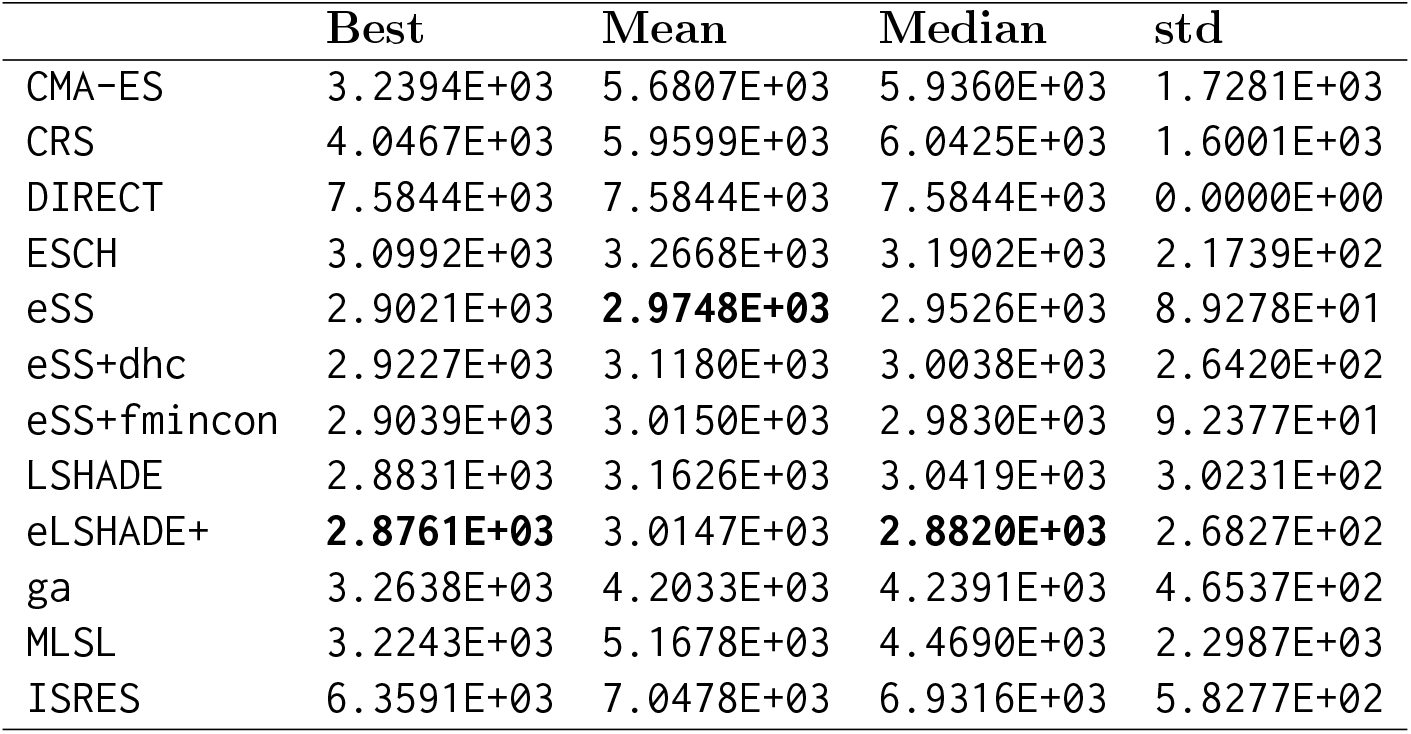
Summary of results for benchmark B5. For each algorithm, we report: the best value of all runs, and the mean, median and standard deviation of all the solutions reported. We highlight: the lowest best solution found, the lowest mean, and the lowest median.

**Table 12:** Summary of results of `LSHADE` and `eLSHADE+` for problems B1_e, B2_e, and B4_e. For each algorithm, we report the best value over all runs, together with the mean, median, and standard deviation of the final solutions. Highlighted values indicate the lowest best, mean, and median for each problem.

| Problem | Algorithm | Best | Mean | Median | std |
| --- | --- | --- | --- | --- | --- |
| B1_e | eLSHADE+ | **1.0168E+04** | **1.6128E+04** | **1.5960E+04** | 3.4976E+03 |
|  | LSHADE | 1.1789E+04 | 2.0767E+04 | 1.9177E+04 | 7.7764E+03 |
| B2_e | eLSHADE+ | **1.3334E+02** | **1.6050E+02** | **1.3512E+02** | 5.0933E+01 |
|  | LSHADE | 1.5392E+02 | 5.5902E+02 | 7.8662E+02 | 2.9960E+02 |
| B4_e | eLSHADE+ | **3.5980E+01** | **3.0486E+03** | **1.0738E+02** | 4.3206E+03 |
|  | LSHADE | 1.1200E+02 | 5.1394E+05 | 6.1337E+03 | 1.0751E+06 |

## 5. Discussion

This section discusses the implications of the results, specifically addressing scalability, the impact of hybridization, and the stability of the convergence behavior.

### 5.1. Scalability and Convergence Stability

The results highlight a clear stratification in algorithmic performance. The only methods that consistently delivered high-quality solutions across all dimensions were `eLSHADE+` (and its variants), the original `LSHADE`, and the `eSS` family. `CMA-ES` was competitive only in benchmark B4, suggesting limited generalizability for this specific class of biological problems.

A critical observation regarding scalability is the temporal behavior of the algorithms. For medium-scale problems, `eSS` typically exhibits faster initial progress. However, `LSHADE` and `eLSHADE+` achieve a more profound reduction in error during the later stages. This behavior is linked to the linear population size reduction mechanism employed by the `DE`-based methods. As the population shrinks, the algorithm naturally shifts from exploration to intensification.

This mechanism also appears to mitigate the phenomenon of finite-time blow-up and other numerical instabilities in the dynamics that some of these dynamic models exhibit for certain regions of the parameter space [109, 110]. These numerical issues introduce sudden, destabilizing spikes in population variance or objective function degradation, which usually damage the performance of the optimizer. As illustrated in Figure 15, the frequency and magnitude of these blow-ups reduce significantly for `DE`-based methods as time advances. The shrinking population acts as a stabilizing force, enforcing convergence and reducing the likelihood of the search diverging into poor regions of the landscape in the final stages.

**Figure 15.**
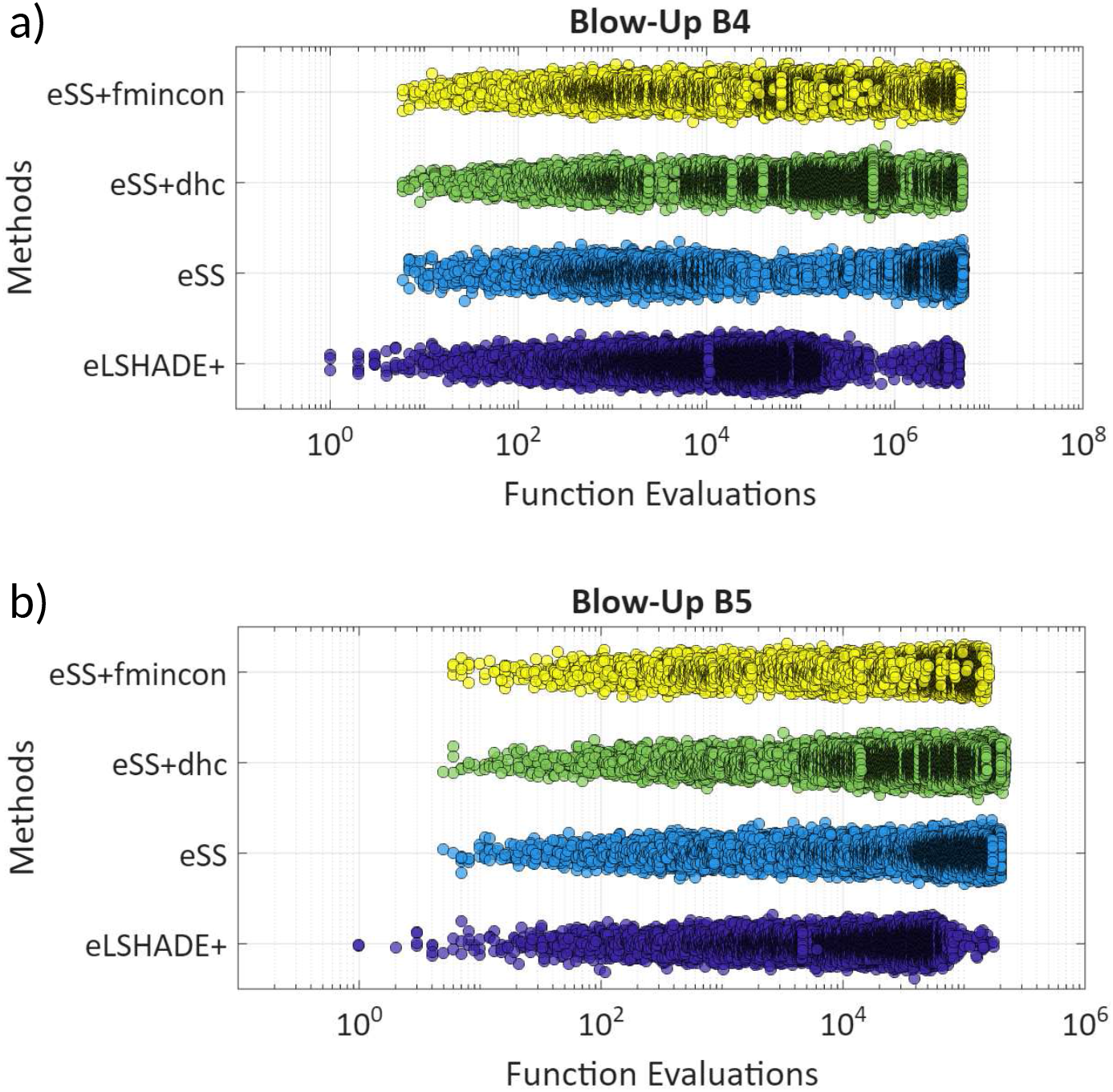
Timeline of numerical blow-ups versus function evaluations for `eLSHADE+`, `eSS` and its hybrid configurations for: a) Benchmark B4, and b) Benchmark B5. Each point represents an objective function evaluation. Vertical clustering indicates numerical instabilities triggering population variance spikes. Note the reduction in blow-up frequency for eLSHADE+ compared to eSS variants, attributed to the population size reduction mechanism

For the large-scale benchmark B1, however, `DE`-based strategies converged faster even in the early stages. This suggests that the adaptive mutation and crossover mechanisms of these methods are inherently better suited to handling the curse of dimensionality than the diversification generation method of scatter search.

Overall, this study represents a change in the state-of-the-art, emerging `eLSHADE+` as the most reliable method to tackle parameter estimation in systems biology-related problems. In fact, it proved not only to be the best algorithm for reducing the objective function value the most, but it also outperformed the other tested algorithms in terms of robustness. It even outmatched `eSS`, which was conceived as the best strategy for this kind of tasks up until now.

### 5.2. The Effect of Hybridization

Our proposed method, `eLSHADE+`, balances exploration and exploitation by integrating periodic local searches. The choice of local optimizer is critical: while gradient-based methods (such as `fmincon`) theoretically offer faster convergence, they can be detrimental if the objective function is noisy or computationally expensive. Conversely, derivative-free methods (`dhc`) are more robust but typically slower.

Our probabilistic selection strategy balances this choice automatically, and the results generally validate this approach. In most benchmarks, `eLSHADE+` achieved better or comparable performance to the standalone `LSHADE`. However, the exception of Benchmark B1, where the `LSHADE` performed slightly better, serves as a reminder that such hybridization is not a panacea. In extremely high-dimensional, expensive landscapes, the computational budget consumed by local search might be better spent on global exploration. Nevertheless, it is worth noting that `eLSHADE+` provides a robust default that performs optimally in the majority of cases.

### 5.3. Synergy of the Multi-Strategy Hybridization

The ablation study confirms that the performance of `eLSHADE+` is not merely the sum of its components, but the result of a synergistic interaction between the adaptive global search and the dual local solvers. As observed in the ablation study section, removing either the gradient-based or the derivative-free component leads to a degradation in performance on specific benchmarks.

This supports the hypothesis that biological landscapes are topologically heterogeneous. For example, in stiff systems like Benchmark B2, the gradient-based solver exploits the smooth valleys created by the fast time-scale dynamics, achieving precision that derivative-free methods struggle to match. Conversely, in the rougher landscapes of Benchmark B5 (Signal Transduction), the derivative-free component prevents the search from stalling in local noise traps where gradient information is unreliable. The probabilistic switching mechanism effectively decouples the optimization logic from the specific defects of the ODE solver, offering a resilience that single-mode hybrids (like standard `eSS`) lack.

Overall, the success of `eLSHADE+` can be attributed to its ability to decouple the decision to intensify from the method of intensification. In some problems, the landscape is riddled with local optima. A rigid hybrid strategy would force a local solver start at every trigger point, consuming evaluations to refine poor solutions. By including a probability of *skipping* local search (the “Pure Exploration” mode), `eLSHADE+` effectively implements a resource-conservation strategy, allowing the population-based mechanism to escape these complicated regions before committing expensive resources to fine-tuning.

## 6. Conclusions

This work introduces `eLSHADE+`, a novel hybrid metaheuristic specifically tailored to address the rigorous demands of parameter estimation in systems biology. Current state-of-the-art hybrids typically rely on rigid, single-mode intensification strategies that struggle with the heterogeneous topology of biological models landscapes characterized by a pathological mixture of stiff, smooth valleys and noisy, rugged plateaus. Our contribution addresses this fundamental limitation through a probabilistic multi-strategy architecture that adaptively decouples the decision to intensify from the method of intensification.

The proposed method enhances the adaptive `LSHADE` framework with two key innovations. First, a probabilistic selection mechanism chooses among three operational modes at each local search trigger: (i) gradient-based search for rapid convergence in smooth valleys, (ii) derivative-free search for robustness against numerical noise and discontinuities, and (iii) pure global exploration that deliberately skips local refinement to preserve computational budget in highly rugged or deceptive regions. This tri-modal hybridization allows the algorithm to adaptively handle both differentiable manifolds and non-smooth regions of the search space. Second, an optional logarithmic transformation addresses the multi-scale nature of biochemical parameters, facilitating efficient exploration when bounds span several orders of magnitude.

Comprehensive benchmarking against the BioPreDyn suite demonstrates that `eLSHADE+` consistently outperforms the previous state-of-the-art method, enhanced Scatter Search (`eSS`), as well as other established algorithms. Across several challenging real-world problems spanning 86 to 1,759 parameters, our method achieves superior solution quality (up to two orders of magnitude improvement on benchmark B3), lower variance across independent runs, and more reliable convergence in late-stage optimization. These results establish `eLSHADE+` as a new baseline for parameter estimation in this domain, particularly for problems involving wide parameter bounds, stiff dynamics, or expensive function evaluations.

However, some limitations and open questions remain. The observed performance variability between structurally similar problems indicates that model topology and computational budget allocation remain critical factors requiring deeper theoretical understanding. The balance of the weighting scheme for probabilistic mode selection may benefit from adaptive strategies that adjust during optimization based on landscape features or convergence metrics. Furthermore, broader evaluation on varied model types, including partial differential equations, stochastic systems, and models with discrete events, will be desirable to fully characterize the method’s applicability.

A particularly promising avenue for future research lies in the development of *cooperative heuristics*. Our analysis revealed distinct temporal performance characteristics: `eSS` often excels in early-stage convergence speed through aggressive diversification, while `eLSHADE+` demonstrates superior late-stage accuracy via its adaptive intensification mechanisms. This complementarity suggests that hybrid frameworks combining both algorithms in parallel, with strategic information exchange (e.g., sharing elite solutions or triggering transitions based on convergence stagnation), could synergistically combine rapid initial exploration with high-precision final refinement. Such cooperative strategies represent a natural evolution of the probabilistic multi-strategy paradigm introduced here.

In conclusion, by systematically addressing the topological heterogeneity of biological objective landscapes through adaptive hybridization, `eLSHADE+` advances the state-of-the-art in robust model calibration for systems biology. The method, along with the integrated evaluation software and comprehensive benchmark data, is publicly available to facilitate reproducible research and further algorithmic development in this challenging domain [25].

## Acknowledgements

J.R.B. acknowledges that this work has received funding from grant PID2023-146275NB-C22 (DYNAMO-bio project) funded by MICIU/AEI/10.13039/501100011033 and ERDF/EU. A.P.R. acknowledges Xunta de Galicia, through the Axencia Galega de Innovación, for a predoctoral grant (Programa de axudas á etapa predoutoral 2025, IN606A-2025/011).

D.R.P. and J.R.B. acknowledge support by the European Commission – NextGenerationEU, through Momentum CSIC Programme: Develop Your Digital Talent. D.R.P. is hired under the Generation D initiative, promoted by Red.es, an organisation attached to the Ministry for Digital Transformation and the Civil Service, for the attraction and retention of talent through grants and training contracts, financed by the Recovery, Transformation and Resilience Plan through the European Union’s Next Generation funds.

## Notes

https://zenodo.org/records/18379328

### Competing Interest Statement

The authors have declared no competing interest.

